# MMP-2/9 inhibition modulates sharp wave abundance, inhibitory proteoglycan sulfation, and fear memory in juvenile zebrafish: relevance to affective disorders

**DOI:** 10.1101/2025.02.25.640105

**Authors:** I. Blanco, S. Deasy, Matthew Amontree, M. Gabriel, A. Caccavano, S. Vicini, E. Glasgow, K Conant

## Abstract

Sharp wave ripple (SWR) events, present in diverse species, spontaneously occur in the hippocampus during quiescent restfulness and slow-wave sleep. SWRs comprise a negative deflection, the sharp wave (SW) event with an often-superimposed ripple (R) and are the neural correlates of memory consolidation and recall. The Anterodorsolateral lobe (ADL) (zebrafish hippocampal homologue) exhibits SW and SWR events, and since SWs initiate SWRs, their abundance typically shows the same directionality. In previous work, we observed matrix metalloproteinase-9 (MMP-9)-dependent effects on depression-relevant behaviors, perineuronal net (PNN) levels, and SWR abundance in the adult rodent hippocampus. Here, we investigate MMP-2/9-dependent effects on biochemical, behavioral, and neurophysiological endpoints in juvenile zebrafish and zebrafish at the transition from the late juvenile period to early adulthood. With MMP-2/9 inhibition, juvenile zebrafish showed reduced SW amplitude and abundance together with increased fear memory retention and reduced sociability. Juvenile zebrafish also showed an increased percentage of longer-duration SW events. Except for a reduction in SW amplitude, these changes were not observed at the transition from late juvenile to early adulthood. These changes were accompanied by increased levels of chondroitin sulfate (CS) proteoglycan 4-*O*-sulfation, which modulates PNNs and excitatory-to-inhibitory (E/I) balance. Discontinuation of MMP-2/9 inhibition in juvenile zebrafish normalized deficits in ADL SW abundance and sociability. Together, these findings show that MMP-2/9 significantly influences E/I balance and learning and memory during the highly plastic juvenile period in zebrafish. Findings also have relevance to an emerging appreciation of PNN changes that may contribute to altered neuronal oscillations and mood or cognition.

## Introduction

Physiological release and regulation of matrix metalloproteinase-9 (MMP-9) contribute to the excitatory to inhibitory (E/I) balance in the developing and mature central nervous system (CNS), which may be crucial to optimal cognition and mental health. Neuronal release of MMP-9 occurs rapidly following *N*-methyl-D-aspartate (NMDA) receptor stimulation of cultured neurons or high-frequency tetanic stimulation of rodent hippocampal slices, and this is followed by dendritic spine expansion (**Szklarczyk et al., 2002; Wang et al., 2008**). Consistent with this, MMP-9 is indispensable for long-term potentiation of synaptic transmission (**Gorkiewicz et al., 2015; Nagy, 2006**). Studies in rodents and zebrafish have also linked MMP-9 activity with increased dendritic arbor (**Alaiyed et al., 2020; Silva et al., 2019**). More recently, MMP-9 has also been shown to reduce brain extracellular matrix (ECM) to potentially increase excitatory neurotransmission. For example, ECM-rich perineuronal nets (PNNs), which predominantly surround fast spiking GABAergic interneurons and increase their activity (**Cabungcal et al., 2013; Frischknecht et al., 2009; Tewari et al., 2018**), are attenuated by MMP-9 activity and their diminution can lead to disinhibition of excitatory neurotransmission. Additionally, secretion of both MMP-2 and −9 from microglia and astrocytes are shown to also remodel PNNs (**Alaiyed et al., 2020; Venturino et al., 2021**). The role of PNNs and their modulators on memory and pyramidal excitability is, however, complicated by regional differences in their expression and the cell types which they surround (**Carstens et al., 2016**). For example, though PNNs generally restrict hippocampal plasticity (**Alaiyed et al., 2019; Carstens et al., 2016; Carulli & Verhaagen, 2021**), they have been shown to terminate infantile amnesia (**Ramsaran et al., 2023**), promote memory retention in brain regions including the amygdala (**Gogolla et al., 2009; Poli et al., 2023**), and to modulate aspects of sociability including social memory (**Carstens et al., 2021; Cope et al., 2022; Huang et al., 2023; Rey et al., 2022**).

PNNs are composed of chondroitin sulfate proteoglycans (CSPGs) containing core proteins such as brevican, aggrecan, versican, and neurocan with sulfated glycosaminoglycan (GAG) side chains with different sulfation patterns including chondroitinin-4-sulfate (CS-4/A) and chondroitin-6-sulfate (CS-6/C) covalently linked to the core proteins (**Meyer-Puttlitz et al., 2002; Ruoslahti, 1996**). Hyaluronic acid and proteoglycan link protein (HAPLNs) associate lecticans at their N-terminal to the hyaluronan acid backbone while tenascins (C and R) cross-link lecticans at their C-terminal. Before the closure of critical periods which in mammals is associated with PNN maturation, PNNs show high levels of CS-6 compared to CS-4-sulfation, and the former are more permissive for PNN proteolysis. In contrast, 4-sulfation is more prevalent in adulthood and decreases the susceptibility of PNNs to proteolysis, thereby rendering them more stable (**Carulli & Verhaagen, 2021; Fawcett & Kwok, 2022; Kitagawa, 2016; Miyata et al., 2012; Miyata & Kitagawa, 2016**). Consistent with this, increasing the CS4/CS6 ratio favors PNN deposition while increasing CS-6-sulfated forms promotes persistent plasticity (**Miyata et al., 2012**). Importantly, proteoglycans, as well as 4-O-and 6-O-sulfotransferases are present during zebrafish development (**Chen et al., 2005; Habicher et al., 2015, 2021**).

In previous work, we observed that a serotonin and norepinephrine reuptake inhibitor, venlafaxine, increased MMP-9 levels in a murine model of depression stimulating an MMP-9-dependent increase in *ex vivo* gamma power and SWR abundance leading to learning and memory improvements (**Alaiyed et al., 2019, 2020; S. et al., 2018**). Importantly, these changes were associated with increased pyramidal cell dendritic arbor and a reduction in PNN levels. Herein we address the question of whether temporal inhibition of MMP-2/9 activity is sufficient to cause specific biochemical, behavioral, and neurophysiological changes that influence mood and/or memory in zebrafish. We further explore the novel question of whether specific effects of MMP-2/9 activity are reversible and conserved across diverse vertebrate species. We chose zebrafish as our model due to its many advantages including their known expression of MMP-2/9 (**Wyatt & Crawford, 2021; Yoong et al., 2007**), their everted brain positioning their hippocampus dorsally (**Folgueira et al., 2012; Ganz et al., 2015**) allowing for non-invasive *in vivo* calcium imaging of neuronal population dynamics through their transparent skin (**Blanco et al., 2024; Mu et al., 2019**), and their hippocampal homologue, the Anterodorsolateral lobe (ADL), which exhibits SWs and SWR events comparable to mammals. Moreover, activity in zebrafish ADL can be measured in whole *ex vivo* brains without damaging integral hippocampal fibers or inputs (**Blanco et al., 2024; Vargas et al., 2012**). Zebrafish also display highly tractable behaviors necessary for studying complex brain disorders (**Bashirzade et al., 2022; Kalueff et al., 2014**).

For the present study, we examined a selective MMP-2/9 inhibitor (SB-3CT), able to cross the blood brain barrier (BBB) (**Meisel & Chang, 2017**), for its effects on select physiological and behavioral endpoints in specific stages of zebrafish development, including early juvenile and the transition from late juvenile to early adulthood. We conducted whole brain *ex vivo* SW recordings from the ADL and we performed behavioral assays to examine locomotion, anxiety/stress, fear memory, and sociability. In addition, we performed whole-brain proteomics to elucidate some of the molecular players potentially contributing to these changes. Since PNN deposition is increased in rodent models of depression and is modulated by knockdown of MMP-9 activity (**Alaiyed et al., 2019, 2020; Riga et al., 2017**) we further investigated telencephalon-specific changes in PNN composition. Additionally, we asked whether neuronal population activity and changes in behavior are reversibly altered by transient inhibition of MMP activity in this model organism.

## Methods

### Experimental Animals

Wild-type early juvenile (30-56 days post-fertilization) and late juvenile/early adult (2-4 months) zebrafish were used, and both age groups (50:50 female:male ratio) were randomly housed in groups of 10-20 in 2L tanks with water temperature kept at 28°C. Zebrafish were fed either once or twice a day and were maintained on a 14/10 light-dark cycle (lights on 9 AM and lights off at 11 PM). Zebrafish were exposed to either 0.002% dimethyl disulfide (DMSO – vehicle) or 3 µM SB-3CT (MMP-2/9 pharmacological inhibitor, Sigma, St. Louis, MO, USA, Cat # S1326) for a period of two consecutive days. Stock solutions for SB-3CT (150 mM) were aliquoted and stored at −20°C until use. The drug was changed daily. After our two consecutive day protocol, independent groups of treated zebrafish underwent either behavior or local field potential (LFP) recordings. Each experiment was independently replicated at least 2 times. Investigators were blinded to treatment groups. All procedures were performed in accordance with the Institutional Animal Care and Use Committee of Georgetown University, Washington DC, USA. Protocol # 2020-0034.

### Local Field Potential Recordings

LFP recordings were performed during the light cycle (between 9 am and 7 pm). Recordings were alternated between treated and non-treated zebrafish and they were obtained from whole-brain *ex vivo* preparations as per our previous study (**Blanco et al., 2024**). Briefly, recordings were obtained using borosilicate glass pipettes filled with artificial cerebrospinal fluid (aCSF) with approximate 150 kΩ tip resistance (Sutter P87) from the Anterodorsolateral lobe (ADL) of carefully extracted whole brains after zebrafish were deeply anesthetized with Tricaine-S (MS-222, Syndel, Bin A02F01G, ANADA # 200-226). Brains were recovered in a chamber filled with oxygenated (95% O_2_, 5% CO_2_) aCSF at high flow rate (20 mL/min) at room temperature. aCSF contained in mM: 134 NaCl, 2.9 KCl, 2.1 CaCl_2_, 1.2 MgCl_2_, 10 HEPES, and 10 Glucose, pH 7.4. Recovery was no less than 30 minutes (**Brenet et al., 2019**). 30-50 minutes of LFP recordings were acquired per animal – each dot in a graph is equivalent to n = 1 zebrafish. Importantly, we observe tissue viability that exceeds 2 hours of reliable recordings above baseline noise. Extracted brains that did not exhibit reliable SW events above baseline noise within 2 minutes of electrode placement (<5%) were excluded from the study.

### Local Field Potential Analysis

Analysis of LFP recordings was performed using a custom MATLAB script previously described (**Caccavano et al., 2020**), with some modifications (**Blanco et al., 2024**). Briefly, 60 Hz line noise and its high-frequency harmonics were removed and a Gaussian finite impulse response band-pass filter with corrected phase delay was applied between 1-1000 Hz and subsequently filtered between 1-30 Hz for detecting the SW and 120-220 Hz for ripples (though the latter was not quantified for this study). The root mean square (RMS) of the SW and SWR was computed in sliding 30 and 5 ms windows, respectively. SW and SWR events were detected from the RMS signals that exceeded 6 and 4 standard deviations (SD) above baseline, while the event start and end times were determined when the RMS signals exceeded 4 and 3 SD above baseline, respectively. Events with a duration of less than 25 ms were discarded, and successive events with less than 150 ms inter-event interval were treated as one continuous event. To determine the baseline SD for each RMS signal in a way that accounts for both inactive and active recordings, the entire time series was binned into a histogram and a two-term Gaussian fitting technique was employed to estimate both the baseline noise and true signal.

### Behavior

All behavioral assays were conducted during the light cycle between 10 am and 7 pm. Immediately prior to beginning each experiment, zebrafish were acclimated to the behavior room for at least 45 minutes. Besides the Novel tank test and Open field paradigms, where Open field directly followed the Novel tank test, all behavioral assays were conducted with independent zebrafish cohorts. Assays were performed in fish water (0.3g sea salt/L) with no drug. Water changes were made consistently throughout the experiments. Locomotion and velocity (Open field test) were analyzed using EthoVision XT 16. The rest of the behavioral assays were analyzed manually with the investigators blinded to the treatment group.

#### Novel tank test

The test was conducted using an 8’ x 4’ x 4’ glass tank with water levels at approximately 3.75’ from the bottom. Each fish was tested individually for five minutes while swimming freely around the arena. A camera facing the front view of the tank recorded the fish’s swimming behavior. Time spent at the top versus the bottom was quantified. The latency to reach the top compartment was also quantified.

#### Open field test

After a five-minute acclimation to the tank (the duration for the Novel tank test), the camera was re-positioned to face the top of the tank. An additional five minutes were recorded while the fish swam freely.

#### Novel Object Location Task (NOLT)

The NOLT test was performed in the same tank as the Open field and Novel tank test. This test consisted of two training days where the fish was allowed to explore the tank for 5 minutes each day. Inside the tank, two identical objects were placed on opposite corners (corners A and B). On the testing day (Day 3), the fish was put back into the same tank but now one of the objects was moved to a novel location (corner C). The time the fish spent exploring the novel versus the familiar locations was quantified. The discrimination index (D.I.) was calculated using the following formula:

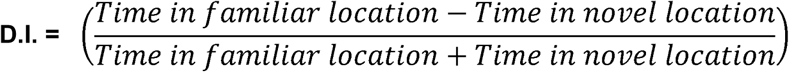

#### Passive avoidance test

This behavioral assay was previously employed by Kim and colleagues **(Kim et al., 2010)** and modified for this study. For this assay, the fish learns to associate an aversive stimulus with crossing from one compartment of the tank to the next. A different tank of similar dimensions as the previous tests, but with a permanent sliding acrylic gate was used. Briefly, each zebrafish was placed into a tank divided by a gate into two different compartments, one with light blue walls and the other with no color (transparent). In our study, we quantified the latency to cross from the light blue compartment (no stimulus) to the transparent one (where the aversive stimulus occurs) as a measurement of the fear response/memory. On Day 1, after a period of two minutes acclimation to the light blue compartment, a cue (lightbox ON for 5 seconds) appears right before the gate is lifted. As soon as the zebrafish crosses compartments, the gate is closed, and a stone at least 3 times the size of the fish (aversive stimulus) is dropped right in front of the fish causing a startle response. Day 1 consisted of four trials: a baseline with no aversive stimulus followed by three consecutive aversive trials. After the last trial, the fish were returned to their home tank for 24 hours to allow for memory consolidation. 24 hours later, the fish was returned to the same arena and after the lightbox was turned ON and the gate was opened, their fear response was measured as the latency to cross compartments.

#### 3-Chamber social test

In this social paradigm, the zebrafish was transferred to a 3 cm thick acrylic plexiglass tank with 8 x 31 x 8 cm dimensions. The tank is divided into three different compartments: the social and object chambers measuring 8 x 12 x 8 cm each and a center compartment measuring 8 x 8 x 8 cm. Both the social and object compartment contain an internal chamber, placed in the middle, measuring 3 x 4 x 8 cm for the placement of a group of 3-4 non-treated age-matched zebrafish, or an inanimate object. The experimental zebrafish was always placed in the center chamber and allowed to freely swim for 10 minutes while a camera (top view) recorded its activity. The time spent in either compartment was manually quantified, and the social preference index was quantified using the following formula:

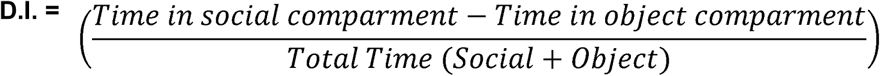

#### Immunofluorescence

Dissected whole brains from deeply anesthetized zebrafish with an overdose of MS-222 were post-fixed in 4% paraformaldehyde (PFA, Thermofisher Scientific, J19943-K2) at 4°C for 48 hours. Brains were cryoprotected in 30% sucrose in phosphate-buffered saline (PBS, Fischer bioreagents, Cat # BP399-500) and kept at 4°C for approximately 72 hours. After sucrose, whole brains were rinsed in PBS and transferred to dry ice for 30 minutes before saving them at −80°C until use.

Whole brains were acclimated to −20°C for at least 30 minutes prior to embedding in optimal cutting temperature (OCT, Fisher HealthCare product Cat # 4585) solution. 20 µm coronal slices were obtained using a Leica CM3050 S Cryostat. Sectioned slices were immediately mounted onto charged microscope slides and stored at 4℃ for up to 2 weeks or −80℃ for long term storage. The day before staining, slices were left at room temperature to acclimate and dry overnight. The following day, slices were baked 30 minutes at 60℃ and a hydrophobic barrier was drawn. Post-fixation with 4% PFA was performed for 1 hour. The slices were washed twice in PBS (2 mins each time) and permeabilized in 0.5% Triton-X (Sigma-Aldrich, X-100, Batch No. 106K0177) in PBS for 90 minutes. Following permeabilization, slices were incubated with blocking buffer: 10% goat serum (Life Technologies, Cat # 50062Z) for 90 minutes. Slices were then washed twice with wash buffer: 0.25% Tween-20 (Millipore Sigma, P2287) in PBS. Primary antibody Anti-Chondroitin Sulfate Antibody (CS-56, Millipore Sigma, SAB4200696) was diluted in 0.2% Tween-20/5% goat serum/ PBS (1:200) and slices incubated overnight at 4℃. Slices were then washed three times (2-3 minutes) and incubated with secondary antibody solution: Alexa Fluor 488 goat anti-mouse IgG (1:1000, ThermoFisher Scientific, Cat # A11001) in 1% goat serum/ PBS for 3 hours. Slices were washed 3 times and cover slipped with Vectashield HardSet™ antifade Mounting Medium with DAPI (Cat # H-1500). Z-stack images were acquired using a Leica SP8 Laser Confocal microscope. Images were processed using Z-project in FIJI.

#### Western blots

Whole-brains lysates from juvenile zebrafish were homogenized in Radioimmunoprecipitation Assay (RIPA) buffer (Thermo Scientific Pierce RIPA buffer, Cat # 89900) with Protease and Phosphatase inhibitor (Thermo Scientific, Cat # 1861281) and centrifuged at 10 000 RPM to discard any debris. Supernatants were transferred to clean tubes and protein quantification was performed using Bio-Rad DC Protein Assay (Cat #s: 500-0114, 500-0113, and 500-0115). 50 µg of total protein was incubated with 20 mU Chondroitinase ABC (ChABC, Sigma-Aldrich, Cat # C367) at 37°C for 2 hours. Lysates were spun down and mixed with Laemmli sample buffer (Bio-Rad, Hercules, CA, USA, Cat # 161-0737) containing 5% β-mercaptoethanol (βME) and subsequently boiled at 95°C for 7 minutes. Control lysates were treated with 0.01% Bovine Serum Albumin (BSA, Vector Laboratories, Cat # SP-5050-500) under the same conditions. Protein separation was achieved by electrophoresis on precast gels (10 or 4-20% mini protein TGX gels, Bio-Rad Cat #s: 4568034 and 4568094, respectively). Proteins were transferred to nitrocellulose membranes (Trans-Blot Turbo Transfer, Bio-Rad, Cat # 1704159) at 2.5A 20V for 14 minutes followed by blocking in 5% non-fat milk in 0.1% Tween-20/ PBS for 1 hour at room temperature. Membranes were washed in 0.1% Tween-20/ PBS for at least 30 minutes with several washes in between. Membranes were probed with primary antibodies overnight at 4°C and with secondary antibodies for 1 hour at room temperature (Supplementary Table S1). Protein bands were visualized using a chemiluminescence machine.

To complement the proteomics results, the telencephalon of 10-15 juvenile zebrafish were removed and pooled for a total of three independent cohorts. Telencephalon were lysed in RIPA buffer containing Halt protease inhibitor Cocktail (Thermo Scientific™, Cat#78438) and total protein was measured with the Pierce™ BCA protein assay (Thermo Scientific™, Cat# 23225). For immunodetection of CSPGs, samples underwent digestion with 0.67 U/ml ChABC for 30 minutes at 37°C. Laemmli buffer containing 5% βME was added to tissue homogenates, incubated at 85 °C for 10 minutes, cooled on ice, and then subjected to electrophoresis on single-percentage SDS-PAGE gels (6 and 10%). Gels were transferred to nitrocellulose membranes (Trans-Blot Turbo Transfer, Bio-Rad, catalog #1704159), blocked for 1 hour in 5% BSA + 0.05% Tween-20 (v/v in TBS) on an orbital shaker at room temperature, and then incubated in the primary antibody overnight at 4°C on an orbital shaker. The next day, membranes were washed 3x for 5-15 minutes with 0.05% Tween-20 (v/v in TBS), incubated for 60-90 minutes in 5% nonfat milk blotting-Grade buffer + 0.05% tween-20 (v/v in TBS, Bio-Rad, Cat# 1706404) containing the appropriate secondary HRP-conjugated antibody (1:5000) on an orbital shaker at room temperature. Membranes were washed 3x for 5-15 minutes with 0.05% Tween-20 (v/v in TBS), washed 2x for 5-15 minutes in TBS, and then subjected to chemiluminescent detection with SuperSignal™ West Pico PLUS Chemiluminescent Substrate (ThermoFisher, Cat# 34579) or SuperSignal™ West Atto Ultimate Sensitivity Substrate (ThermoFisher, Cat# 38554). The West Atto substrate was diluted at 1:15 in West Pico PLUS when West Pico PLUS demonstrated insufficient chemiluminescent detection. Membranes were stripped with restore western blotting stripping buffer (Thermo Scientific™, Cat# 21059) for 15 minutes at 37°C followed by 15 minutes at room temperature on the orbital shaker. Membranes were washed 3x for 10-15 minutes in 0.05% Tween-20 (v/v in TBS) prior to incubation with the subsequent primary antibody (Supplementary Table S1).

### nanoUPLC-MS/MS

Protein samples were prepared in the same manner as those for whole-brain lysate Western blot. An average of 6 zebrafish whole brains were pooled for each n (n = 5 DMSO/SB-3CT-treated). 100 µg of total protein was sent to the GUMC-Proteomics Shared Resources for the proteomics analysis using the Orbitrap Lumos Tribrid mass spectrometer instrument.

#### Sample preparation

Proteins in pellets were extracted by using lysis buffer containing 10% SDS, 1x proteinase inhibitor cocktail and 50 mM triethylammonium bicarbonate. After the addition of benzonase, the cellular suspension was incubated on ice for 20 min followed by sonication with a probe-tip sonicator for 5 pulse (10 sec on 20 sec off for each pulse). The cell lysate was centrifuged for 15 min, 16000 g at 4 °C, with the supernatant saved and used for protein concentration determination by BCA assay. Equal amount of proteins from each sample was then treated with DTT and iodoacetamide and then loaded onto a S-Trap column (ProtiFi, LLC) by following the manufacturer’s instructions. Proteins were digested with sequencing-grade Lys-C/trypsin (Promega) by incubation at 37°C overnight. The resulting peptides were eluted and dried down with a SpeedVac (Fisher Scientific). To construct a comprehensive spectral library, an aliquot of each digest was pooled and fractionated by high pH reversed phase fractionation into 8 fractions, peptides in each fraction were dried down with a SpeedVac.

#### NanoUPLC-MS/MS

Peptides are analyzed with a nanoAcquity UPLC system (Waters) coupled with Orbitrap Fusion Lumos mass spectrometer (Thermo Fisher). Samples in 0.1% FA solution are loaded onto a C18 Trap column (Waters Acquity UPLC M-Class Trap, Symmetry C18, 100 Å, 5 μm, 180 μm x 20 mm) at 10 µL/min for 4 min. Peptides are then separated with an analytical column (Waters Acquity UPLC M-Class, peptide BEH C18 column, 300 Å, 1.7 μm, 75 μm x 150 mm) with the temperature controlled at 40°C. The flow rate is set as 400 nL/min. A 150-min gradient of buffer A (2% ACN, 0.1% formic acid) and buffer B (0.1% formic acid in ACN) is used for separation: 1% buffer B at 0 min, 5% buffer B at 1 min, 22% buffer B at 90 min, 50% buffer B at 100min, 98% buffer B at 120 min, 98% buffer B at 130 min, 1% buffer B at 130.1 min, and 1% buffer B at 150 min. Data files were acquired with data independent acquisition (DIA) mode or data dependent acquisition (DDA) on an Orbitap Fusion Lumos mass spectrometer using an ion spray voltage of 2.4 kV and an ion transfer temperature of 275°C. Mass spectra are recorded with Xcalibur 4.0. Advanced peak determination is on for MS analyses. MS parameters for data dependent acquisition (DDA) mode were set as below: Detector Type: Orbitrap; Mass range: 375-1500 m/z; Orbitrap Resolution: 120,000; Scan Range: 375-1500 m/z; RF Lens: 30%; AGC Target: Standard; Maximum Injection Time Mode: Auto; Microscans: 1. Charge state: 2-9 s; Exclusion duration: 40 s; Cycle Time: 3 s. MS/MS parameters were set as below: Isolation Mode: Quadrupole; Isolation Window: 1.6 m/z; HCD normalized collision energy: 35%; Detector Type: Orbitrap; Resolution: 30,000; Normalized AGC Target: 200%. MS parameters for data independent acquisition (DIA) mode: MSⁿ Level (n):2; Isolation Mode: Quadrupole; Activation Type: HCD; HCD Collision Energy (%): 28; Detector Type: Orbitrap; Orbitrap Resolution: 30000; Mass Range: Normal; Scan Range Mode: Define m/z range; Scan Range (m/z): 200-1800; RF Lens (%):30; AGC Target: Custom; Maximum Injection Time Mode: Dynamic; Desired minimum points across the peak: 6; Microscans: 1.

#### Data analysis

Analysis of DDA raw files was performed in Proteome Discoverer (Thermo Fisher Scientific, version 2.4) with Sequest HT database search engines. The database with Zebrafish was downloaded from Uniprot. The database-searching parameters were set as below: full tryptic digestion and allowed up to two missed cleavages, the precursor mass tolerance was set at 10 ppm, whereas the fragment-mass tolerance was set at 0.02 Da. Carbamidomethylation of cysteines (+57.0215 Da) was set as a fixed modification, and variable modifications of methionine oxidation (+15.9949 Da), acetyl (N-terminus, +42.011 Da) were allowed. The false-discovery rate (FDR) was determined by using a target-decoy search strategy. The decoy-sequence database contains each sequence in reverse orientations, enabling FDR estimation. On the peptide level, the corresponding FDR was less than 1%. Analysis of DIA raw files was done by using Spectronaut (Biognosys, v15) with hybrid library built from 8 DDA data files (from high pH reversed phase fractionation) and 10 DIA data files. All settings are default. In brief, dynamic retention time prediction with local regression calibration was selected. Interference correction on MS and MS2 level was enabled. The Qvalue Cutoff was set to 1% at peptide precursor and protein level using scrambled decoy generation and dynamic size at 0.1 fraction of library size. MS2-based quantification by area was used, enabling local cross-run normalization.

#### Statistics

All statistical tests were done using GraphPad Prism 9. Data normality and outlier test were performed using the Shapiro-Wilk test and the ROUT outlier test, respectively. Tukey’s multiple comparison test, Šídák’s multiple comparisons test, and Uncorrected Fisher’s LSD were conducted as *post-hoc* tests following a significant 2Way ANOVA with an alpha value of > 0.05 when appropriate. Unpaired Student’s t test and Whitney Mann tests were also performed when looking at 2 groups with data that passed the normality test or not, respectively. Paired Student’s t-test and the Chi-square were also performed when appropriate.

Figures were created using Biorender

## Results

### MMP-2/9 inhibition decreases exploration and locomotion in an anxiety/stress-independent manner

Systemic MMP-2/9 inhibition was achieved by exposing zebrafish to the pharmacological inhibitor SB-3CT, previously shown to cross the BBB (**Meisel & Chang, 2017**), in their water. SB-3CT was not toxic to the fish following a two-day protocol of 3 µM exposure as evidenced by >96% survival rates (data not shown). We first investigated whether MMP-2/9 inhibition affected locomotion in early juvenile and late juvenile/early adult zebrafish (Fig. 1A-C). We saw a significant decrease in the total distance traveled in the Open field test in early juvenile (N_DMSO_ = 21, N_SB-3CT_ = 17; Mann-Whitney U = 80.50, p = 0.0033) as well as a decrease in their mean velocity (N_DMSO_ = 21, N_SB-3CT_ = 17; Mann-Whitney U = 75, p = 0.0019) (Fig. 1B). The decrease in total distance traveled (N_DMSO_ = 19, N_SB-3CT_ = 18; Unpaired t_(35)_ = 3.271, p = 0.0024) and mean velocity (N_DMSO_ = 19, N_SB-3CT_ = 18; Unpaired t_(35)_ = 3.310, p = 0.0022) was also evident in late juvenile/early adult zebrafish (Fig. 1C).

**Fig. 1:**
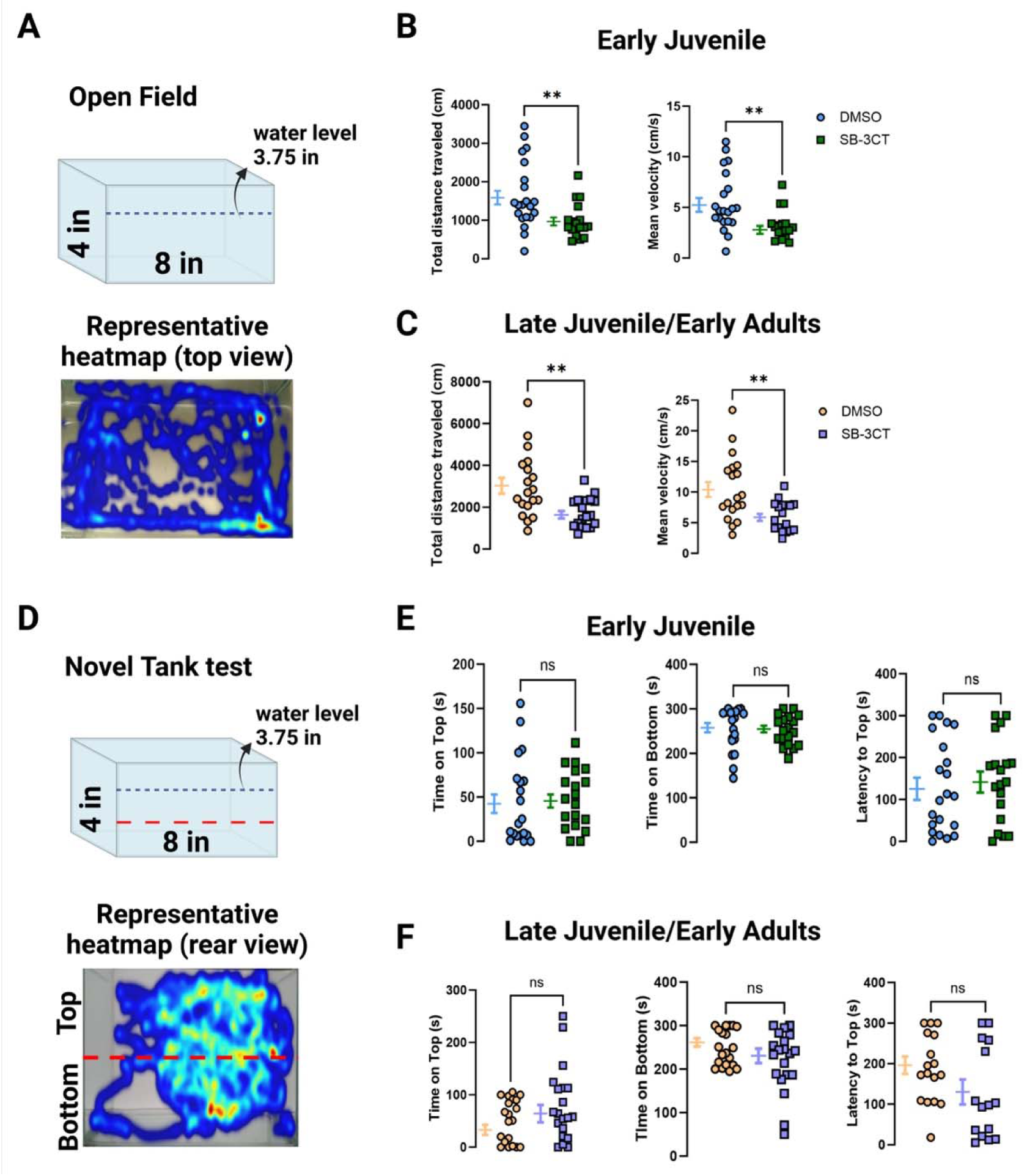
MMP-2/9 inhibition decreases locomotion independently from anxiety/stress in zebrafish. **A** Top panel: representative cartoon of the Open field tank with dimensions and water level. Bottom panel: representative heatmap of the explorative trajectory of a juvenile DMSO-treated zebrafish. **B** Quantification of total distance traveled and mean velocity for early juvenile zebrafish. **C** Quantification of total distance traveled and mean velocity for late juvenile/early adult zebrafish. **D** Top panel: representative cartoon of the Novel Tank with dimensions, water level, and division of top versus bottom compartments (red dashed line). Note that the same setup was used to measure both behaviors. Bottom panel: representative heatmap of the trajectory of a juvenile non-treated zebrafish showing their time in each compartment. **E** Quantification of the time spent on either top or bottom compartment in the Novel Tank test as well as latency to reach the top of the tank for early juvenile zebrafish. **F** Quantification of the same parameters for late juvenile/early adult zebrafish. Unpaired Student’s t and Mann-Whitney tests. Graphs are expressed as mean ± S.E.M. Data were acquired with EthoVision XT16 software. **p < 0.01.

To rule out the possibility that a reduction in locomotion was due to anxiety or stress, we quantified data from the Novel tank test (Fig. 1D-F). The time spent on the top compartment (N_DMSO_ = 20, N_SB-3CT_ = 19; Mann-Whitney U = 162, p = 0.4395), bottom compartment (N_DMSO_ = 20, N_SB-3CT_ = 19; Mann-Whitney U = 162, p = 0.4397), or the latency to reach the top compartment (N_DMSO_ = 21, N_SB-3CT_ = 19; Mann-Whitney U = 181.5, p = 0.6341) were not different between groups in early juvenile zebrafish. Similar results were found for late juvenile/early adults with no differences in the time spent on the top (N_DMSO_ = 20, N_SB-3CT_ = 20; Mann-Whitney U = 154, p = 0.2176), bottom compartment (N_DMSO_ = 20, N_SB-3CT_ = 20; Mann-Whitney U = 154, p = 0.2176), or the latency to reach the top compartment (N_DMSO_ = 17, N_SB-3CT_ = 15; Mann-Whitney U = 78, p = 0.0618) between groups (Fig. 1D-F). Though SB-3CT inhibits both MMP-2/9, the overall decrease in locomotion may be linked to MMP-9 inhibition, as this has been observed in the rodent literature (**Hernandez-Anzaldo et al., 2016**), and this effect appears to be independent of both developmental stage and anxiety/stress.

### MMP-2/9 inhibition increases spatial retention of familiar location in early juvenile zebrafish

Multiple studies support the involvement of MMP-9 in long-term memory consolidation in mammals (**Gorkiewicz et al., 2015; Nagy, 2006; Wright & Harding, 2009**). Therefore, to examine whether MMP-9 activity is required for long-term memory consolidation in the zebrafish, we performed the NOLT test (Supplementary Fig. S1A). In this assay, zebrafish are trained to learn the spatial location of two identical objects for two consecutive days, and then on the third day one of the objects is moved to a novel location (corner C). We found that inhibiting MMP-2/9 activity in early juvenile zebrafish increased the bias towards spending majority of time in the familiar quadrant (corner B) during recall when compared to the non-treated group (N_DMSO_ = 13, N_SB-3CT_ = 13; Unpaired t_(24)_ = 2.299, p = 0.0305) (Supplementary Fig. S1B). Contrary to this finding, inhibiting MMP-2/9 activity in late juvenile/early adult zebrafish did not bias them toward the familiar quadrant (N_DMSO_ = 12, N_SB-3CT_ = 12; Unpaired t_(22)_ = 1.404, p = 0.1743) (Supplementary Fig. S1C). Together, the data may suggest that MMP-9 and possibly MMP-2 inhibition differentially affect long-term spatial object recognition memory in early juvenile zebrafish. The data could further suggest an increased fear of novelty in juvenile zebrafish that is absent at the transition to adulthood.

### MMP-2/9 inhibition enhances fear retention in early juvenile zebrafish

Based on the possibility of increased fear of novelty in the NOLT and the role of MMP-9 in fear memory in mammals (**Bach et al., 2018; Grochecki et al., 2023; Knapska & Kaczmarek, 2016**), we wondered if MMP-9’s role in modulating fear response and memory was also present in zebrafish. To test this, we adapted the passive avoidance assay from Kim and colleagues (**Kim et al., 2010**) (Fig. 2A). Briefly, a zebrafish is placed in one of two differently colored compartments divided by a physical barrier (acrylic door) for 2 minutes after which the barrier is lifted, and an aversive stimulus (the drop of a stone) is presented as soon as the zebrafish crosses compartments. After three trials, an aversive response is created to the compartment where the aversive stimulus is presented, in this case the transparent compartment. The zebrafish learn to associate their behavior of crossing to the transparent compartment with a punishment (aversive stimulus) and avoid crossing to that compartment. The aversive compartment was chosen after preliminary data showed that zebrafish had no preference for either compartment/color (data not shown) and that their latency to cross to either compartment under non-stressful circumstances was under 100 seconds. To signal the opening of the acrylic door, a lightbox placed under the tank was turned ON for 5 seconds before the gate opened. The transparent compartment was better suited to letting the light through as opposed to the light blue bottom of the blue compartment. As soon as the zebrafish crossed compartments, the gate was closed, and the stone (aversive stimulus) was dropped in front of the zebrafish. After the aversive stimulus was presented, the zebrafish were immediately (∼5 seconds) returned to their home tank. On Day 1, this process was performed three consecutive times while the latency to cross from the blue to the transparent compartment was recorded. To measure the fear response retention, the zebrafish were returned to the same arena 24 hours after the last trial on Day 1, and the latency to cross from the blue to the transparent compartment after the light turned ON and the gate opened was measured. An increase in the latency to cross to the transparent compartment was interpreted as an indication of the fear memory retention.

**Fig. 2:**
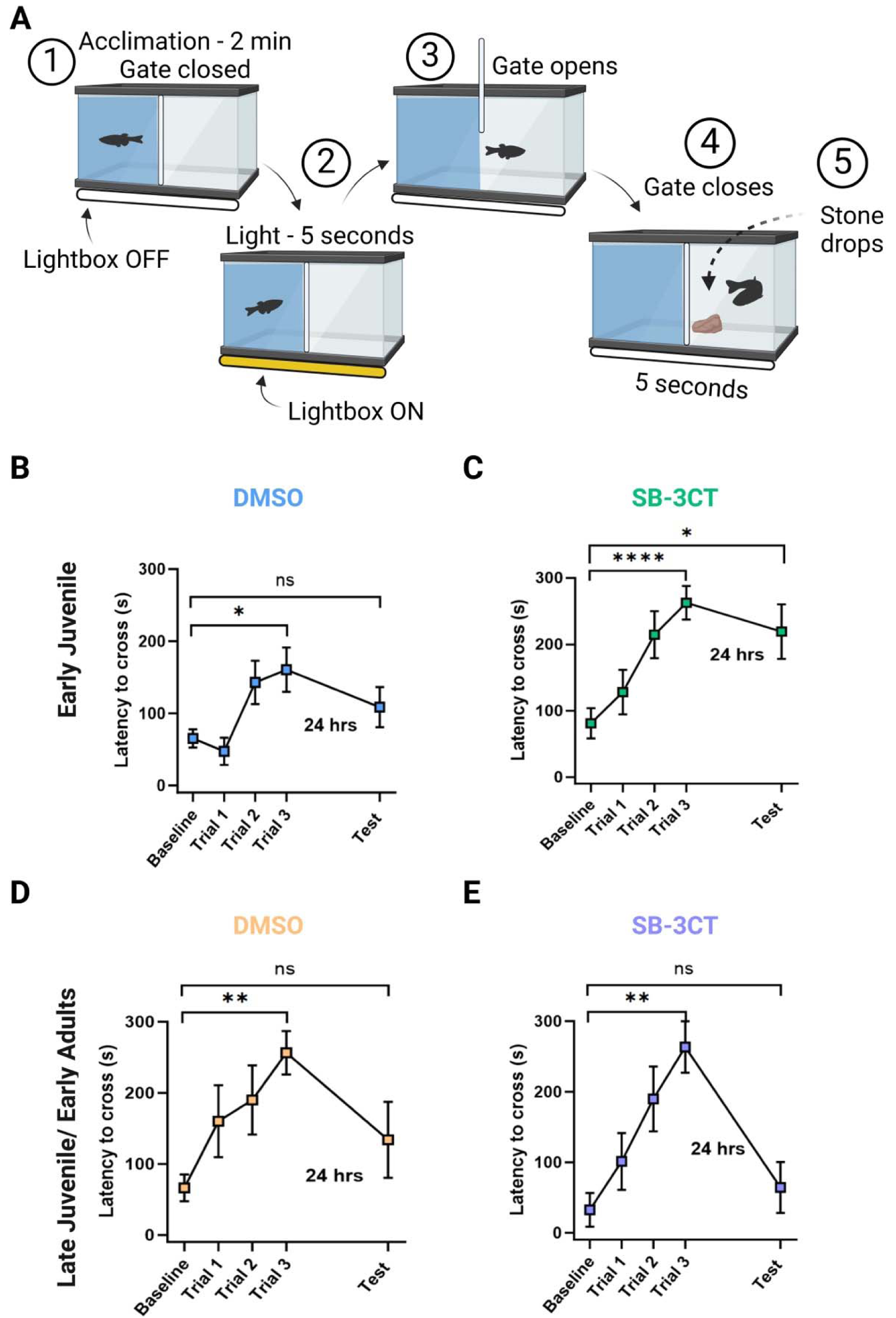
MMP-2/9 inhibition enhances fear memory retention in early juvenile zebrafish. **A** Representative cartoon of the Passive avoidance behavior setup. Numbers correspond to: (1) acclimation, (2) lightbox ON, (3) gate opens, (4) gate closes, (5) stone drops. **B–E** Graphs represent the latency to cross from the blue compartment to the transparent compartment at baseline and three consecutive trials on Day 1, and 24 hours later (Day 2) for **B–C** DSMO- and SB-3CT-treated early juvenile zebrafish and **D–E** DSMO- and SB-3CT-treated late juvenile/early adult zebrafish. Data is represented as mean ± S.E.M. Independent Paired Student’s t tests were performed between baseline and trials on Day 1 and Mann-Whitney tests were performed between baseline and testing day. *p < 0.05, **p < 0.01, ****p < 0.0001.

As expected, the average baseline latency to cross compartments for both early juvenile (N_DMSO_ = 15-16, N_SB-3CT_ = 10-12) and late juvenile/early adult (N_DMSO_ = 7, N_SB-3CT_ = 7) zebrafish whether treated or not, was under 100 seconds (Fig. 2B-E). By trial 3 of Day 1, DMSO-treated zebrafish had learned to associate their behavior of crossing compartments with ‘punishment’ as shown by a significant increase in their latency to cross to the transparent compartment (baseline: 65.06 ± 12.68 secs vs. trial 3: 160.50 ± 30.82 secs; Paired t_(14)_ = 2.663, p = 0.0177) (Fig. 2B). However, 24 hours later, the fear response was abolished as evidenced by a non-significant difference in the latency to cross to the transparent compartment between baseline (65.06 ± 12.68 secs) and the testing day (108.50 ± 27.86 secs; Mann-Whitney U = 105.5, p = 0.5783) (Fig. 2B). Similarly, after MMP-2/9 inhibition, there was a significant difference in the latency to cross to the transparent compartment between baseline (81.00 ± 22.70 secs) and the third trial (262.58 ± 25.34 secs) in early juvenile zebrafish (Paired t_(11)_ = 6.461, p < 0.0001), suggesting learning of the fear response (Fig. 2C). However, unlike control zebrafish the fear memory persisted 24 hours later in the SB-3CT-treated group, as evidenced by a sustained increase in the latency to cross to the transparent compartment, where the aversive stimulus was previously presented, from baseline (81.00 ± 22.70 secs) to testing (219.20 ± 41.27 secs; Mann-Whitney U = 24, p = 0.0153) (Fig. 2C). Interestingly, the difference between groups by the third trial (DMSO: 160.50 ± 30.82; SB-3CT: 262.58 ± 25.34 secs) was also significant, (Mann-Whitney U = 51, p = 0.0206), which could reflect more rapid encoding/learning of the fear memory after MMP-2/9 inhibition.

We then tested SB-3CT treated and non-treated late juvenile/early adult zebrafish (Fig. 2D-E). DMSO-treated zebrafish showed a significant difference in the latency to cross compartments from baseline (66.43 ± 18.96 secs) to the third trial on Day 1 (256.43 ± 30.55 secs; Paired t_(6)_ = 5.074, p = 0.0023) (Fig. 2D). However, they were not able to retain the association between the aversive stimulus and crossing to the transparent compartment 24 hours later (baseline latency: 66.43 ± 18.96 vs. testing day latency: 134.14 ± 53.68 secs; Mann-Whitney U = 23, p = 0.8695). Similarly, after MMP-2/9 inhibition, late juvenile/early adult zebrafish learned to associate crossing compartments with ‘punishment’ as evidenced by a significant difference in the latency to cross from baseline (32.57 ± 24.19 secs) to the third trial (263.43 ± 36.57 secs; Paired t_(6)_ = 5.130, p = 0.0022). However, unlike the effect of MMP-2/9 inhibition on early juvenile zebrafish, MMP-2/9 inhibition did not aid in the retention of the fear response 24 hours later in late juvenile/early adults (baseline latency: 32.51 ± 24.19 secs vs. testing day latency: 64.29 ± 36.16 secs; Mann-Whitney U = 16, p = 0.3036) (Fig. 2E). No significant difference was found between groups during the third trial (Mann-Whitney U = 19, p = 0.5594), suggesting that encoding of the fear memory was unaffected by MMP-2/9 inhibition in late juvenile/early adults. Statistical results are shown in Supplementary Table S2. Together, the data corroborate the differential effect of MMP-9 activity and possibly MMP-2, on zebrafish fear learning and memory retention as a function of age (youth versus the transition into adulthood).

### MMP-2/9 inhibition decreases sociability in early juvenile zebrafish

Studies have shown the potential role of MMP-9 activity in modulating sociability and social anxiety (**Pirbhoy et al., 2020; Reinhard et al., 2015**). Given that early juvenile zebrafish had a fear response that was greater than control and persistent, we next wondered if this increased in fear response would generalize to other domains such as social anxiety/reduced sociability. To measure sociability and the willingness to seek out social settings in this highly social organism; we employed the 3-Chamber social test (Fig. 3A). In this test, zebrafish are exposed to three equal-sized chambers divided into the Social (containing other zebrafish of equal size and age), the Center (where the experimental zebrafish is first placed at the start of the experiment), and the Object (where an inanimate object was placed). For five minutes, the experimental zebrafish are free to swim between chambers, and the time spent in each compartment is measured.

**Fig. 3:**
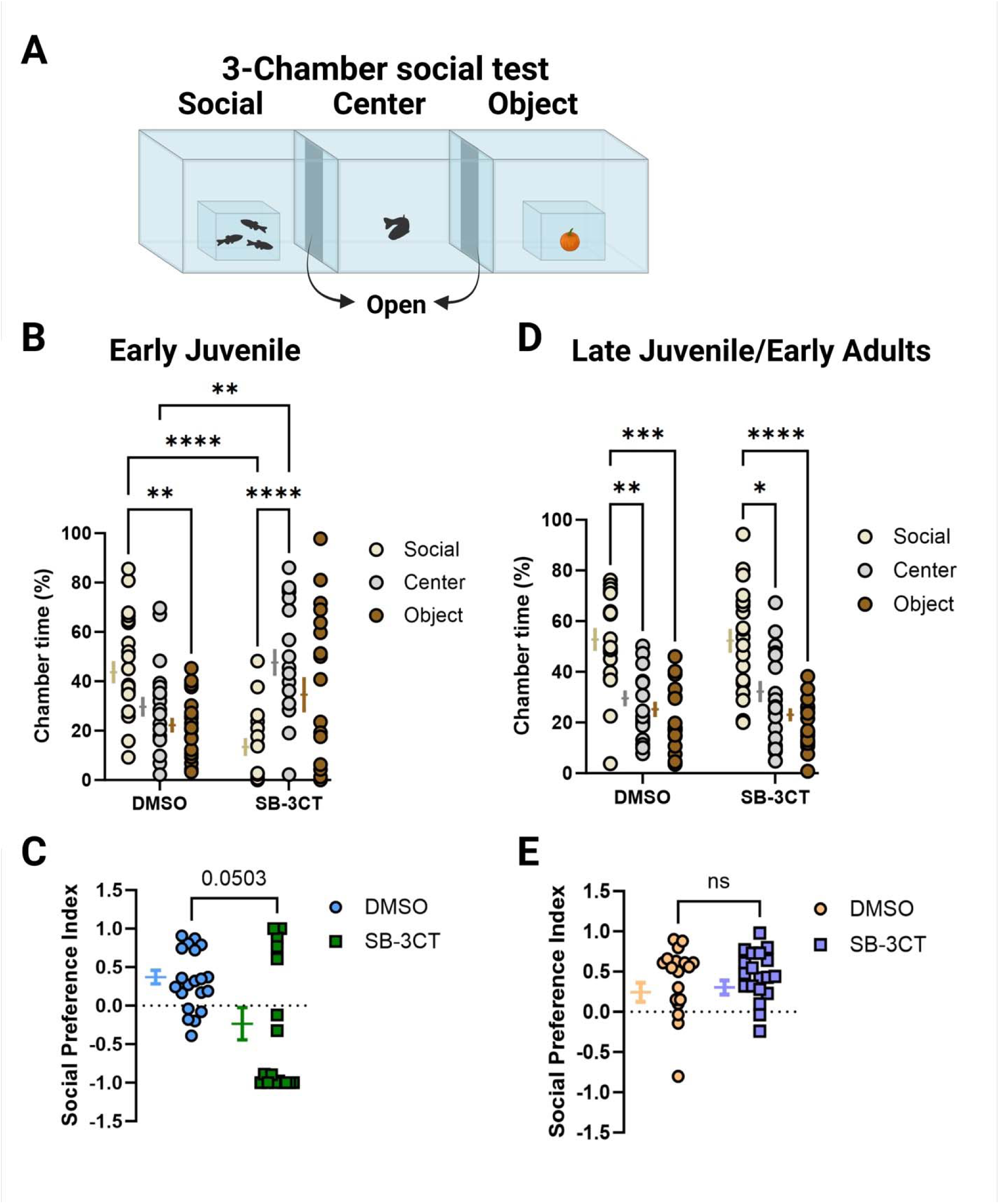
MMP-2/9 inhibition decreases sociability in early juvenile zebrafish. **A** Representative cartoon of the 3-Chamber social test. **B** Shows the percentage of time spent in either chamber for early juvenile zebrafish treated with either DMSO (left) or SB-3CT (right). 2Way ANOVA with Tukey’s multiple comparisons. **C** Shows the social preference index for DMSO- and SB-3CT-treated early juvenile zebrafish. Mann-Whitney test. **D** and **E** are representations of the same measurements but for late juvenile/early adult zebrafish. Graphs are represented as mean ± S.E.M. *p < 0.05, **p < 0.01, ***p <0.001, ****p < 0.0001

A significant interaction between groups and chambers was found in early juvenile zebrafish (F_(2, 72)_ = 13.85, p < 0.0001). As expected, control early juvenile zebrafish (N_DMSO_ = 20) spent significantly more time in the social chamber compared to the object chamber (p = 0.0025). In contrast, inhibiting MMP-2/9 activity significantly shifted the preference of early juvenile zebrafish (N_SB-3CT_ = 18) away from the social chamber toward the center one (p < 0.0001) (Fig. 3B). A trend toward a significant decrease in the social preference index between groups (DMSO: 0.3193 ± 0.0864, SB-3CT: −0.2753 ± 0.2064) was also observed in early juvenile zebrafish (Mann-Whitney U = 113, p = 0.0503) (Fig. 3C). For late juvenile/early adults, no significant interaction was found between groups and chambers (F_(2, 72)_ = 0.2111, p = 0.8102). However, there was a significant difference in time spent in each chamber (F_(1.404, 50.54)_ = 30.31, p < 0.0001), where control zebrafish (N_DMSO_ = 19) preferred the social chamber significantly more than the center (p = 0.0028) and object chamber (p = 0.0005) (Fig. 3D). In contrast to early juvenile treated zebrafish, late juvenile/early adult treated zebrafish (N_SB-3CT_ = 19) did not show any shift in their social preference but rather it mirrored the control group (social vs. center: p = 0.0341; social vs. object: p = < 0.0001). Furthermore, no difference was found in the social preference index between control (0.4007 ± 0.0956) and treated (0.4497 ± 0.0702) late juvenile/early adult zebrafish (Mann-Whitney U = 179, p = 0.9712) (Fig. 3E). Together, the data suggest that MMP-2/9 inhibition differentially affects sociability for early juvenile zebrafish.

### MMP-2/9 inhibition reduces the abundance of SW events in early juvenile zebrafish

To better understand how MMP-2/9 inhibition could differentially affect long-term memory in zebrafish based on their developmental stages, we performed LFP recordings from their hippocampus homologue: ADL (Fig. 4) (**Blanco et al., 2024; Vargas et al., 2012**). Long-term memory consolidation relies on hippocampal-dependent spontaneous events named SWR events that happen during slow wave sleep, quiescent periods, and in *ex vivo* preparations (**Buzsáki, 2015**). As the result of the interplay between pyramidal and parvalbumin (PV)+ GABAergic neurons, an increase in SW events may reflect an increased excitation to inhibition (E/I) balance in the hippocampus and therefore can be modulated by changes in MMP-9 activity (**Alaiyed et al., 2020**). Given that in juvenile zebrafish, a small portion (∼9%) of sharp wave (SW) events contain an embedded ripple (SWR) event (**Blanco et al., 2024**), and that the abundance of SW and SWRs typically follows the same directionality, we focused on MMP-2/9- modulation of SW events, also crucial for learning and memory (**Battaglia et al., 2004; Born, 2010; Shein-Idelson et al., 2016**). Consistent with previously mentioned reports showing that MMP-9 activity modulates the E/I balance in the brain, we found that MMP-2/9 inhibition significantly decreased the abundance (N_DMSO_ = 29, N_SB-3CT_ = 29; DMSO: 22.59 ± 1.686 vs. SB-3CT: 13.70 ± 0.7745/min; Mann-Whitney, U = 163, p < 0.0001) (Fig. 4C) and amplitude (N_DMSO_ = 22, N_SB-3CT_ = 27; DMSO: 5.544 ± 0.6861 vs. SB-3CT: 3.693 ± 0.4323 µV; Mann-Whitney, U = 193, p = 0.0362) (Fig. 4D) of SW events in early juvenile zebrafish. Interestingly, this was also accompanied by an increase in the average duration of these events (N_DMSO_ = 29, N_SB-3CT_ = 29; DMSO: 310.9 ± 18.60 vs. SB-3CT: 429.1 ± 23.16 ms) (Mann-Whitney, U = 188, p = 0.0002) (Fig. 4E).

**Fig. 4:**
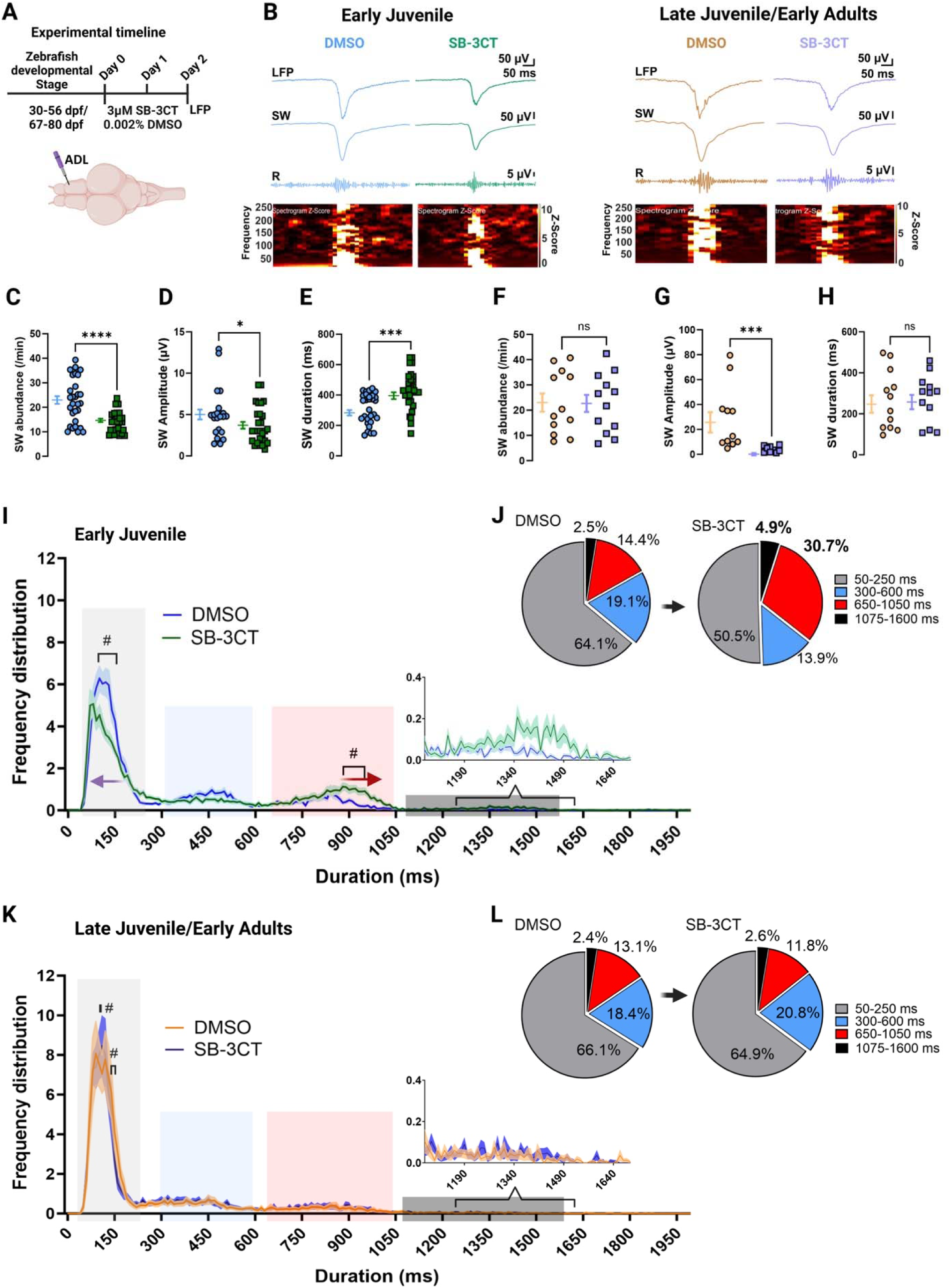
MMP-2/9 inhibition decreases the abundance and amplitude of SWs while enriching the percentage of longer duration SW events in early juvenile zebrafish. **A** Top: Experimental timeline for SB-3CT drug exposure and LFP recordings. Bottom: representative cartoon image of electrode placement in the ADL of *ex vivo* whole brain preparations. **B** Representative traces of SWR events from early juvenile zebrafish treated and not treated with SB-3CT (left) and late juvenile/early adults (right). LFP recordings (1-1000 Hz), filtered SW events (1-30 Hz), filtered ripple events (120-220 Hz), and frequency spectrogram. **C–E** Quantification of SW abundance, amplitude, and duration between DMSO- and SB-3CT-treated early juvenile zebrafish. **F–H** Quantification of SW abundance, amplitude, and duration between DMSO- and SB-3CT-treated late juvenile/early adult zebrafish. Mann-Whitney and Unpaired t tests. **I** Frequency distribution of SW duration for early juvenile zebrafish and **K** for late juvenile/early adult zebrafish between groups. Duration clusters are color-coded for identification: light gray represents cluster 1 (50-250 ms), light blue represents cluster 2 (300-600 ms), light red represents cluster 3 (650-1050 ms), and dark gray represents cluster 4 (1075-1600 ms). Solid lines correspond to the mean and the shaded area shows the S.E.M. The purple arrow shows a shift to the left in the frequency distribution of SW events from treated zebrafish whereas the red arrow shows a shift to the right. 2way ANOVA with Šídák’s multiple comparisons test (# represents significance). **J,L** Quantification of the percentage of SW events within a specified cluster and their change after MMP-2/9 inhibition. Graphs are expressed as mean ± S.E.M. *p < 0.05, ***p < 0.001, ****p < 0.0001.

These results differed from what we observed in late juvenile/early adult zebrafish. There was not a significant change in the abundance (N_DMSO_ = 12, N_SB-3CT_ = 12; DMSO: 23.05 ± 3.612 vs. SB-3CT: 22.71 ± 3.3378/min; Unpaired t, t_(22)_ = 0.0689, p = 0.9457) (Fig. 4F) or duration (N_DMSO_ = 12, N_SB-3CT_ = 12; DMSO: 266.6 ± 41.52 vs. SB-3CT: 276.4 ± 33.70 ms; Unpaired t, t_(22)_ = 0.1820, p = 0.8573) (Fig. 4H) of SW events after MMP-2/9 inhibition. However, MMP-2/9 inhibition significantly decreased the amplitude of these events (N_DMSO_ = 11, N_SB-3CT_ = 9; DMSO: 28.47 ± 7.755 vs. SB-3CT: 4.141 ± 0.8305 µV; Mann-Whitney, U = 4, p = 0.0001) (Fig. 4G), consistent with a decrease in the population of principal cells contributing to each event and a potential reduction in E/I balance, across developmental stages.

An important physiological property of SWR events is their duration, which in mammals ranges between 50 and 150 ms (**Buzsáki, 2015; Vargas et al., 2012**) with a small proportion of longer duration events. Longer duration events are increased during demanding mnemonic tasks and are thought to facilitate the transfer of more relevant information possibly via increasing the number of embedded ripple events (**Davidson et al., 2009; Fernández-Ruiz et al., 2019**). We thus analyzed the frequency distribution of SW duration after SB-3CT treatment using 10 ms non-overlapping time bins. Our data corroborate previous studies showing that most SW events lie between 50 and 150 ms (**Buzsáki, 2015; Vargas et al., 2012**) for all tested groups (events were pooled from all brain recordings: Early juvenile: N_DMSO_ = 22, 230, N_SB-3CT_ = 15, 117; and late juvenile/early adults: N_DMSO_ = 8, 721, N_SB-3CT_ = 8, 382) (Fig. 4I-L). Interestingly, the frequency distribution revealed the existence of four distinct clusters for both groups in the ranges of 50-250, 300-600, 650-1050, and 1075-1600 ms (Fig. 4I, K – clusters are color-coded). Strikingly, in early juvenile zebrafish, MMP-2/9 inhibition reduced the number of shorter duration events (50-250 ms, purple arrow) while enriching the percentage of longer duration (650-1050 ms) SW events (red arrow) (Fig. 4I). We then pooled all the SW events per brain/group and quantified the percentage change of events for each identified cluster after MMP-2/9 inhibition (Fig. 4J, L). In SB-3CT treated early juvenile zebrafish, the percentage of events between 50-250 ms were decreased (0.8-fold) and those between 300-600 ms were also decreased (0.7-fold), while the percentage of events between 650-1050 and 1075-1600 ms were increased (2.1-fold and 2-fold, respectively; Fig. 4J). A significant difference was found between groups when comparing the pooled number of events in the 50-250 ms and 650-1050 ms range after MMP-2/9 inhibition (Chi-square, L^2^ = 1410.41, p < 0.0001). Pooled absolute values corresponding to each duration cluster for all groups are shown in Supplementary Table S3. This pattern was not observed in late juvenile/early adults (Fig. 4K-L). Furthermore, no significant difference was found in the pooled absolute event number between cluster 1 and 3 (Chi-square, L^2^ = 3.54, p = 0.0600). Together, the data suggest that MMP-2/9 inhibition reduces the overall abundance and amplitude of SW events in early juvenile zebrafish while also increasing the percentage of longer duration SW events.

### Effects of MMP-2/9 inhibition on SW events, locomotion, and sociability in early juvenile zebrafish are reversible

We then investigated whether the effects of SB-3CT on SW events and behavior, particularly sociability, in early juvenile zebrafish were transient and dependent on constant exposure. After our two-day inhibition protocol, zebrafish were returned to different tanks without DMSO or SB-3CT for either 5 or 9 days (Fig. 5A). Five days after (d.a.) ‘disinhibition’, there was a persistent decrease in SW event abundance in treated zebrafish (N_DMSO_ = 8, N_SB-3CT_ = 8; Uncorrected Fisher’s LSD, p = 0.0033) that was abolished by day 9 (N_DMSO_ = 8, N_SB-3CT_ = 8; Uncorrected Fisher’s LSD, p = 0.7449) (Fig. 5B). Additionally, 5 d.a. treatment, the average SW duration was not different between groups (N_DMSO_ = 8, N_SB-3CT_ = 8; Uncorrected Fisher’s LSD, p = 0.4363) and we also saw this at 9 d.a. treatment (N_DMSO_ = 8, N_SB-3CT_ = 8; Uncorrected Fisher’s LSD, p = 0.5901) (Fig. 5C). The frequency distribution of the duration of SW events 9 d.a. treatment corroborated that most SW events were between 50 and 200 ms (pooled events: N_DMSO_ = 3, 426, N_SB-3CT_ = 3, 322) (Fig. 5D). Though there was a significant decrease in the number of SW events in this range post-SB-3CT, no significant difference was found between groups when comparing the pooled number of events in the 50-250 ms (N_post-DMSO_ = 1, 811, N_post-SB-3CT_ = 1, 643) and 650-1050 ms (N_post-DMSO_ = 1, 034, N_post-SB-3CT_ = 978) range after MMP-2/9 inhibition (Chi-square, L^2^ = 0.5513, p = 0.4578). In the Open field, post-treated zebrafish showed no significant difference in distance traveled (N_post-DMSO_ = 15, N_post-SB-3CT_ = 17; Mann-Whitney U = 85, p = 0.1136) or mean velocity (N_post-DMSO_ = 15, N_post-SB-3CT_ = 17; Mann-Whitney U = 84, p = 0.1050) (Fig. 5E). We further conducted the 3-Chamber social test 9 d.a. treatment and compared the results with those obtained while the zebrafish were exposed to SB-3CT (Fig. 5F). There was a significant interaction between groups and day of experiment (F_(1, 72)_ = 16.59, p = 0.0001). However, no significant difference in the time spent in the social chamber was found between groups post-exposure (N_DMSO_ = 19, N_SB-3CT_ = 18; p = 0.7632). Notably, we saw a significant increase in sociability for both groups post-exposure compared to groups during DMSO/SB-3CT exposure. Together, these data suggest that MMP-2/9 activity differentially and reversibly modulates SW events, locomotion, and sociability in early juvenile zebrafish.

**Fig. 5:**
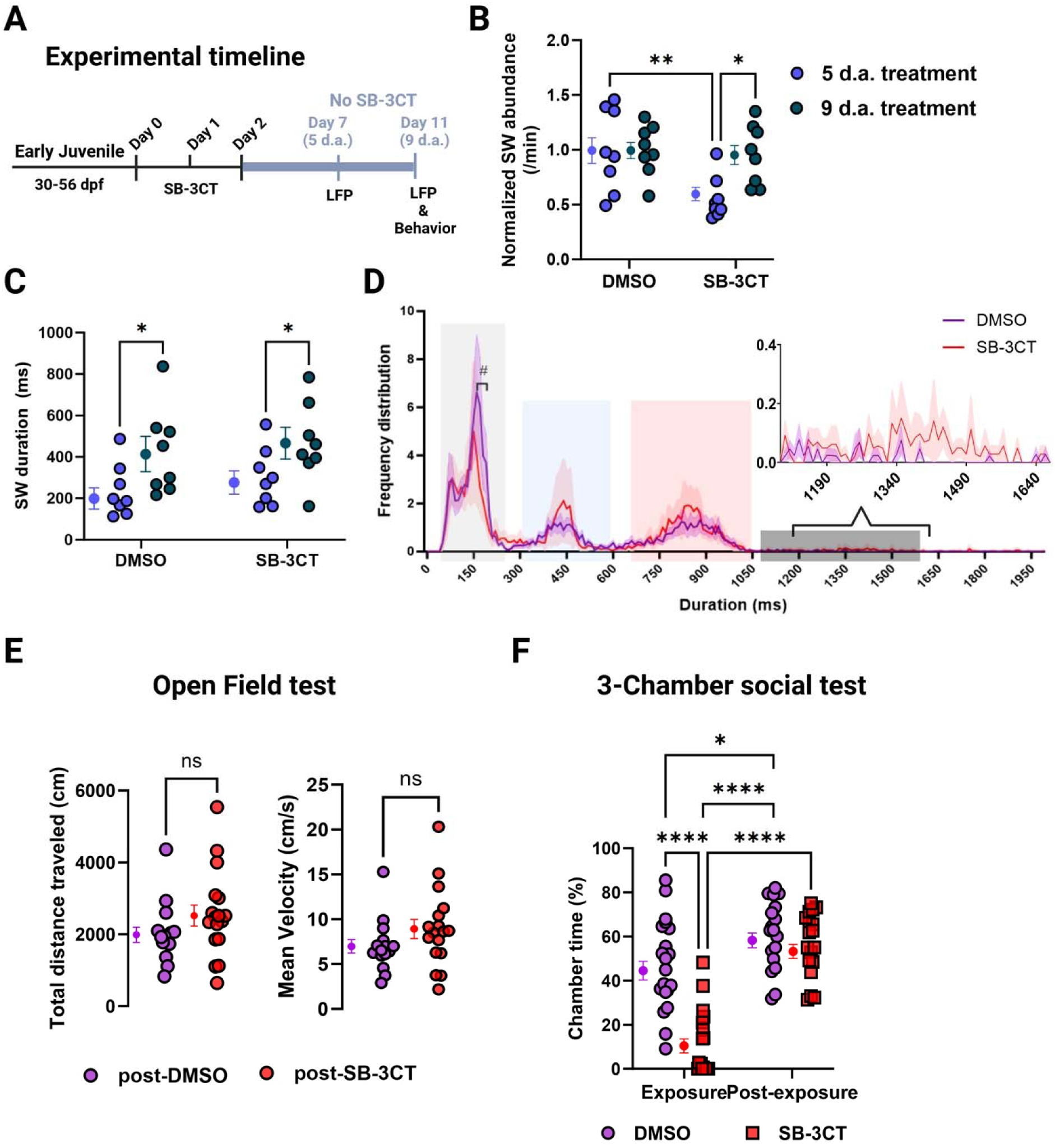
MMP-2/9 ‘dis-inhibition’ reverses changes in SW events and behavior in early juvenile zebrafish. **A** Time line of experiments. **B** Normalized quantification of SW abundance 5 and 9 d.a. DMSO or SB-3CT treatment. 2way ANOVA with Uncorrected Fisher’s LSD. **C** Quantification of the average duration of SW events between groups at 5 and 9 d.a. 2way ANOVA with Uncorrected Fisher’s LSD. **D** Frequency distribution of SW duration for early juvenile zebrafish 9 d.a. treatment with either DMSO or SB-3CT. Duration clusters are color-coded for identification: light gray represents cluster 1 (50-250 ms), light blue represents cluster 2 (300-600 ms), light red represents cluster 3 (650-1050 ms), and dark gray represents cluster 4 (1075-1600 ms). Solid lines correspond to the mean and the shaded area shows the S.E.M. 2way ANOVA with Šídák’s multiple comparisons test (# represents significance). **E** Quantification of total distance traveled and mean velocity. Data were acquired with EthoVision XT16 software. Mann-Whitney test. **F** Quantification of sociability during exposure (from Fig. 4B) and post-exposure. 2way ANOVA with Tukey’s multiple comparison’s test. Graphs are expressed as mean ± S.E.M. *p < 0.05, ****p < 0.0001.

### MMP-2/9 inhibition differentially modulates the proteome of early juvenile versus late juvenile/early adult zebrafish: A potential role for 4-*O*-sulfated CSPGs

Given the importance of PNN modulation and composition in learning and memory and sociability (**Alaiyed et al., 2020; Cope et al., 2022; Fawcett et al., 2022; Huang et al., 2023; Miyata et al., 2012; Miyata & Kitagawa, 2016; Sun et al., 2018**), we searched for PNN-like structures in the telencephalon of zebrafish by immunostaining. We used the pan marker for chondroitin sulfate (CS-56) antibody raised against sulfated CSPGs, which are highly expressed in the hippocampus during development (**Irvine & Kwok, 2018; Takiguchi et al., 2021**). Unlike CS-56, the widely used *Wisteria Floribunda Agglutinin* (WFA) staining mainly recognizes non-sulfated CSPGs and has failed to stain PNNs that are aggrecan positive (**Belliveau et al., 2024; Härtig et al., 2022**), an important PNN component (**Sanchez et al., 2023**). Additionally, CS-56 allows for a more comprehensive search of PNNs across developmental stages. Consistent with previous studies (**Takeda et al., 2018; Zhang et al., 2022**), we observed CS-56+ PNN-like structures throughout the telencephalon of both early juvenile (Supplementary Fig. S2) and late juvenile/early adult (Supplementary Fig. S3) zebrafish, that were mostly located in the dorsolateral region (Fig. 6A – early juvenile), where the ADL is located.

**Fig. 6:**
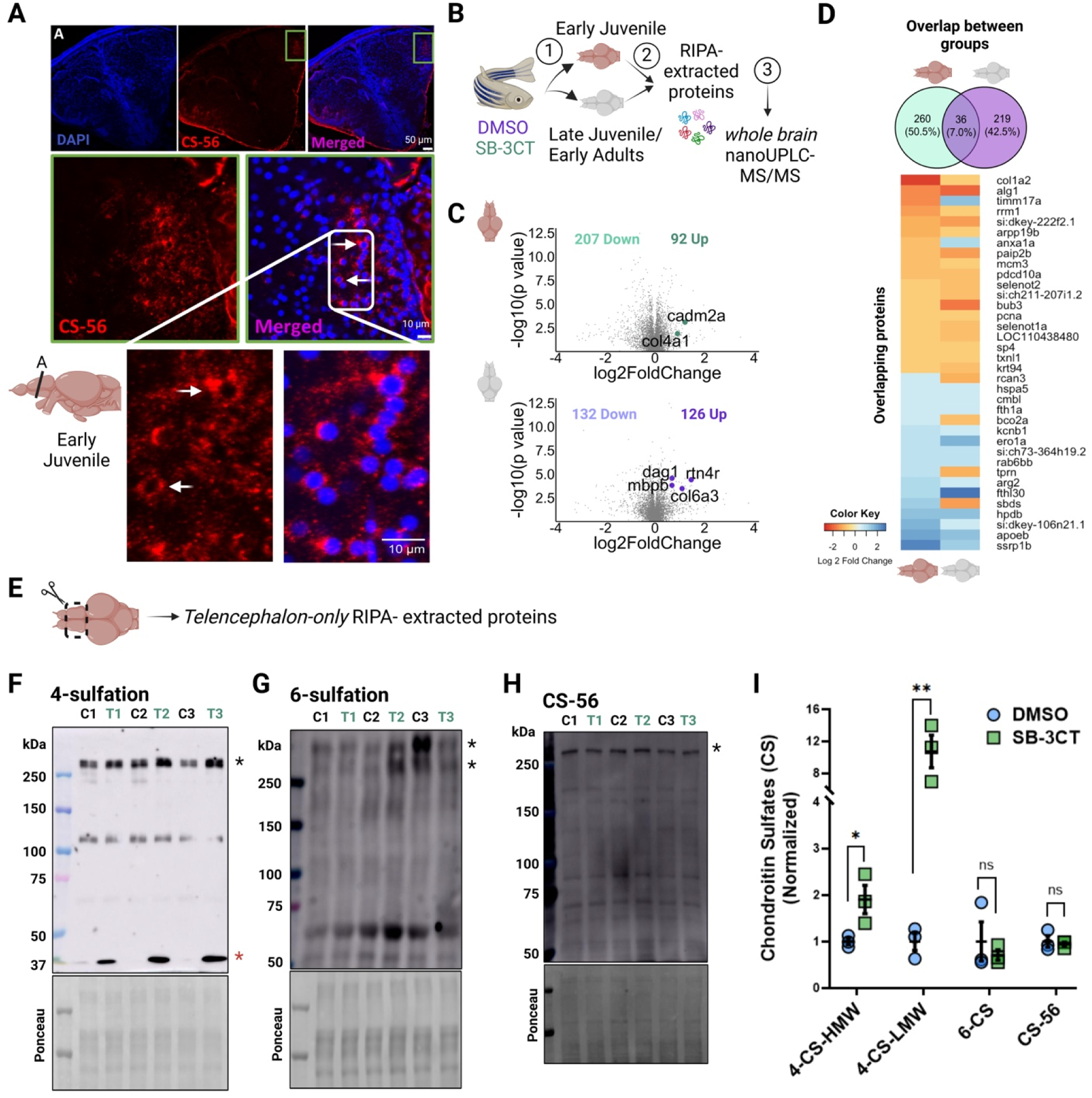
MMP-2/9 inhibition increases plasticity restricting 4-*O*-sulfated CSPGs with no difference in plasticity permissive 6-*O*-sulfated CSPGs. **A** Representative image of chondroitin sulfate (CS)-rich PNNs surrounding the cell body of cells (white arrows) within the telencephalon of early juvenile zebrafish. **B** Representative workflow of the proteomics assay using whole brain lysates. **C** Volcano plots showing downregulated and upregulated proteins after MMP-2/9 inhibition with identified known MMP-9 substrates in both groups. **D** Shows the percentage of overlapping and non-overlapping proteins between groups. **E** Representative cartoon image of a zebrafish brain showing the part of the brain (telencephalon-only) used for Western blot analyses. **F-H** Representative pooled telencephalon-only Western blots for 4-*O*-sulfation, 6-*O*-sulfation, and CS-56, respectively. An average of 10-15 zebrafish telencephalon-only were pooled per group (n=3). For 4- and 6-*O*-sulfation, samples were first digested with ChABC. **I** Quantification of Western blots. Quantified bands are highlighted using asterisks. Specifically, for 4-O-sulfation high molecular weight (HMW) band is denoted by the black asterisk while the low molecular weight (LMW) band is denoted by the red asterisk. For 6-*O*-sulfation quantification, both HMW bands were averaged for quantification. Graphs are expressed as mean ± S.E.M. Unpaired t-test. *p < 0.05, ** p < 0.01.

We then conducted whole-brain nano UPLC-MS/MS (Fig. 6B) to parse out molecular players that could be modulated by MMP-2/9 inhibition affecting E/I balance and learning and memory in early juvenile zebrafish compared to late juvenile/early adult zebrafish. In early juvenile zebrafish MMP-2/9 inhibition significantly downregulated 207 while upregulating 92 proteins. In late juvenile/early adults, MMP-2/9 inhibition significantly downregulated 132 and upregulated 126 proteins (Fig. 6C). A 7% overlap of these proteins was found between groups (Fig. 6D). A full list of significantly changed proteins for both groups detected by our MS analysis are uploaded to Dryad: https://datadryad.org/stash/share/ZbQ1dTCoMfgKieozTZL-wi_1bExY8LVUf5d9Oqx6_eA.

Consistent with previous reports on MMP-9 substrates, SB-3CT significantly increased cell adhesion molecule levels including cell adhesion molecule 2a (cadm2a, fold change 1.210, p = 0.0008) in early juvenile fish and dystroglycan (dag1, fold change 0.6832, p < 0.0001) in late juvenile/early adult zebrafish (Fig. 6C) (**Burmeister et al., 2023; Michaluk et al., 2007**). Cleavage of cell adhesion molecule ectodomains, also part of the ECM, are known to affect neuronal plasticity (**Conant et al., 2015**). Components of the ECM such as collagen, an additional MMP-9 substrate (**Burmeister et al., 2023; LeBert et al., 2015; Mondal et al., 2020**), were also significantly modulated. For example, the MMP-9 substrate collagen 4 (col4a1) was increased in early juvenile zebrafish (fold change 0.9052, p = 0.0133) and collagen 6 (col6a3) in late juvenile/early adult zebrafish (fold change 1.0898, p = 0.0003) (Fig. 6C).

Interestingly, though no core component of the PNN made our Qvalue cutoff of 1%, we saw an increase in versican when this PNN component was evaluated independently (n = 5, Unpaired Student’s t test, t_(8)_ = 2.850, p = 0.0215), as well as a trend for both brevican (t_(8)_ = 2.215, p = 0.0576) and neurocan (t_(8)_ = 2.166, p = 0.0622) in early juvenile zebrafish. No changes in aggrecan b were seen (t_(8)_ = 0.3324, p = 0.7481). Contrary to early juvenile zebrafish, no significant or trending changes were observed in versican (t_(8)_ = 1.770, p = 0.1146), brevican (t_(8)_ = 0.4141, p = 0.6897), neurocan (t_(8)_ = 0.9199, p = 0.3845), or aggrecan b (t_(8)_ = 1.285, p = 0.2348) in late juvenile/early adult zebrafish (Data not shown).

Though proteomics is relatively quantitative and unbiased, a crucial limitation of our analysis is that it was conducted using whole brain lysates. Furthermore, this proteomic analysis does not differentiate between the intact and cleaved lectican species, nor does it address important post-translational modifications. In complementary studies with a separate cohort of control and SB-3CT-treated early juvenile zebrafish, we thus isolated the telencephalon of zebrafish containing their hippocampal homologue (Fig. 6E) and performed Western blot for different PNN components as well as sulfation patterns. Before running telencephalon-only samples, we confirmed that CS proteoglycan immunoreactivity is modulated by ChABC, which unmasks binding sites for the proteoglycan antibodies, in whole brain lysates (data not shown). Telencephalon-only Western blots did not detect statistically significant differences in the expression levels of aggrecan, brevican, neurocan, or HAPLNs 1, 2 or 4 between groups (Supplementary Fig. S4). Our data suggest that SB-3CT does not alter the overall expression of CSPGs or HAPLNs. Western blots conducted to detect specific sulfation forms of CSPGs, however, showed a significant increase in the abundance of 4-sulfated proteoglycans with the MMP inhibitor. Specifically, SB-3CT was associated with a significant increase in high and low molecular weight 4-sulfated CSPGs species (4-CS-LMW: DMSO: 1.00 ± 0.1916 vs. SB-3CT: 10.71 ± 2.049, Unpaired t_(4)_ = 4.720, p = 0.0092; 4-CS-HMW: DMSO: 1.00 ± 0.0710 vs. SB-3CT: 1.901 ± 0.3040, Unpaired t_(4)_ = 2.886, p = 0.0448) (Fig. 6F, I). Additionally, there were no changes in the 6-sulfated CSPGs (average of both bands: DMSO: 1.00 ± 0.4182 vs. SB-3CT: 0.7102 ± 0.1116, Unpaired t_(4)_ = 0.6696, p = 0.5398) (Fig. 6G, I). Similarly, no changes were observed in CS-56 antibody detected CSPGs (DMSO: 1.00 ± 0.1259 vs. SB-3CT: 0.9401 ± 0.0451, Unpaired t_(4)_ = 0.4479, p = 0.6774) (Fig. 6H, I). Though ADL-specific histological analysis would help to determine whether the 4-sulfated CSPGs are increased in PNN structures with SB-3CT treatment, our data suggest a shift in the sulfation pattern of CSPGs in early juvenile zebrafish with MMP-2/9 inhibition. The differential increase in 4-sulfation has previously been shown to make PNNs less susceptible to cleavage, more inhibitory, and to increase PV+ neuronal maturation (**Miyata et al., 2012**), thereby increasing overall inhibition.

## Discussion

In the present study, we show that MMP-2/9 inhibition affects biochemical, behavioral, and neurophysiological endpoints, important to E/I balance, memory, and sociability, in a manner that is more profound in early juvenile zebrafish than during the transition from late juvenile to early adulthood. Specifically, behavioral and physiological changes in early juvenile zebrafish included reduced locomotion, enhanced fear memory retention, decreased sociability, and decreased E/I balance, endpoints which may be altered in association with mood disorders. These changes were further associated with increased levels of ECM components and most importantly with a significant increase in 4-*O*-sulfated CSPGs.

Reduced locomotion has been described in MMP-9 knockout mice (**Hernandez-Anzaldo et al., 2016**), and is observed in a mouse strain with relatively reduced levels of MMP-9 and increased susceptibility to depression-related behavior (**Kennedy-Wood et al., 2021**). Locomotion may also be reduced in humans with major depressive disorder (**Schuch et al., 2017**). Depression-like behaviors are also associated with altered fear memory (**Coutellier & Usdin, 2011**) and PNN deposition (**Banerjee et al., 2017; Fawcett et al., 2022; Gogolla et al., 2009; Hylin et al., 2013; Jovasevic et al., 2021; Ruzicka et al., 2022**). In this study, we found enhanced fear memory retention that was specific to early juvenile zebrafish exposed to SB-3CT. In mammals, fear memory retention is dependent on the integrity of amygdala-specific PNNs (**Gogolla et al., 2009; Poli et al., 2023**). In the zebrafish, we found the presence of CS-56+ PNN structures throughout the telencephalon, where the amygdala is also located, and following MMP-2/9 inhibition, telencephalon-only lysates from early juvenile zebrafish showed increased levels of 4-*O*-sultafation. 4-*O*-sulfation has been associated with decreased PNN turnover as they shift PNNs into a mature phenotype (**Miyata et al., 2012**) that may result in the protection of fear memories. This would be consistent with the rodent literature where the presence of PNNs stabilize fear memory (**Carulli et al., 2010; Gogolla et al., 2009; Poli et al., 2023**). Moreover, though PNNs predominantly surround PV expressing inhibitory interneurons in several brain regions, they are shown to also surround excitatory neurons in the amygdala where they can increase excitatory neuronal activity to increase fear memory (**Morikawa et al., 2017**), a potential mechanism that could contribute to the increase in fear memory retention in early juvenile zebrafish after SB-3CT-induced increase in 4-*O*-sulfated CSPGs.

In addition to enhancing fear memory, changes in PNN integrity could also drive the decrease in sociability seen in early juvenile zebrafish during SB-3CT treatment. Indeed, studies have shown that social dysfunction, including decreased sociability and impaired social memory are, in part, due to changes in PNN deposition in the hippocampal *cornu ammonis* (CA)2 region of rodents (**Carstens et al., 2021; Cope et al., 2022; Domínguez et al., 2019; Huang et al., 2023; Mattioni et al., 2024; Rey et al., 2022**). Normalization of CA2 PNNs was shown to reverse these impairments (**Carstens et al., 2021; Cope et al., 2022; Huang et al., 2023; Rey et al., 2022**). Most recently, it was shown that ablating 4-*O*-sulfation in adult mice within the CA2 disrupted PNN integrity and resulted in reversible mood and social memory impairments (**Huang et al., 2023**). Additionally, though not directly quantified in this study, we hypothesize that early juvenile zebrafish exhibit increased fear in response to social novelty (**Koyuncu et al., 2019; Mattioni et al., 2024**). Moreover, SB-3CT treated early juvenile zebrafish showed improved familiar object location recognition that could reflect the cognitive stability/inflexibility that is thought to be enhanced by PNN maturation (**Fawcett & Kwok, 2022; Happel et al., 2014**), shown to correlate with increased levels of 4-*O*-sulfated CSPGs and associated with major depressive disorder (**Stange et al., 2016**). In contrast to human and rodent studies, however, in which depressive states and/or increased PNN levels generally increase anxiety-related behaviors (**Alaiyed et al., 2019, 2020; Riga et al., 2017**), we did not observe an effect of MMP-2/9 inhibition on anxiety/stress in zebrafish. A caveat, however, is that this data is based only on the Novel tank test. Therefore, a more in-depth study should be conducted to thoroughly investigate the role of MMP-2/9 inhibition on depression, stress, and anxiety-like behaviors in zebrafish, a model organism proposed to be an invaluable tool to study depression (**Nguyen et al., 2014**). Particularly depression should be studied at the juvenile period when according to our data, zebrafish are most vulnerable to MMP-2/9-dependent plasticity.

While biochemical and behavioral endpoints examined in this study show some overlap with human and rodent studies, we observe a relatively more robust conservation of neuronal population changes after MMP-2/9 inhibition. A shift in the E/I balance in the hippocampus is observed as measured by an overall reduction in the abundance of SW events in early juvenile zebrafish as well as a reduction in SW amplitude in both groups. These endpoints could follow from reduced excitability of pyramidal cells in terms of the frequency with which neuronal assemblies are activated, and/or or the population size of activated assemblies (amplitude). As previously suggested, reduced excitability of pyramidal cells could in turn follow from the ability of MMP-9 inhibition to modulate PNNs surrounding PV+ interneurons and thus PV+ mediated inhibition (**Alaiyed et al., 2020; Sun et al., 2018**). Indeed, though from whole brain lysates, which warrants further ADL-specific studies, MMP-2/9 inhibition affected voltage-gated potassium channel Kv3.1b, previously found to be an organizer of both excitatory and inhibitory heterologous synapses and involved in the proper maintenance of neural circuits (**Lee et al., 2017; Pelkey et al., 2015**).

We also found an increase in the percentage of longer-duration SW events in SB-3CT-treated early juvenile zebrafish. This shift was specific to exposure to SB-3CT as recordings during post-exposure did not demonstrate this shift. Therefore, it will be appropriate for future studies to investigate whether these longer duration SW events carry more information important to learning and memory after MMP-2/9 inhibition and whether the enrichment in longer duration SW events are associated with increased trains of ripples including ripple doublets, which have been observed in zebrafish (**Blanco et al., 2024**) and are more likely to occur with longer SWs. Indeed, it would be important to determine if, in association with reducing the abundance of SW events, MMP-2/9 inhibition also increases longer duration events with higher reactivation and replay (**Giri et al., 2024**). Together, our data suggest that, as is the case in rodent models, MMP-2/9 activity likely enhances overall excitatory transmission in the hippocampus/hippocampal homologue.

As compared to early juvenile zebrafish, effects of MMP-2/9 inhibition are relatively subtle in late juvenile/early adult zebrafish. This could reflect physiological changes of MMP-9 levels during development versus the transition to adulthood. Similar to mammals, zebrafish MMP-9 levels are increased during development and during the start of specific developmental stages (**Yoong et al., 2007**). In adulthood, MMP-9 increases may instead be more inducible occurring with neuronal activity and/or microglial activation (**LeBert et al., 2015; Silva et al., 2019**). Due to developmental differences in MMP-9 activity, it is tempting to speculate that PNNs during the early juvenile period are more susceptible to MMP-9 proteolytic cleavage and/or other downstream proteases to help maintain a highly plastic environment conducive to critical periods. In this manner, inhibition of MMP-9 activity during the early juvenile period would modulate PNN deposition and PV+ neurons to potentially decrease overall excitation and concomitantly decrease the abundance of SW events (**Tewari et al., 2018**). In contrast, in a more mature system, with decreased levels of MMP-9 activity, PNNs would be less susceptible to MMP-9 mediated proteolytic cleavage. Consistent with this are previous studies showing that PNNs composed of 6-sulfated CSPGs are more permissive to proteolysis and are highly expressed during mammalian development, whereas 4-sulfated CSPGs are more prevalent in adulthood and significantly decrease the susceptibility of PNNs to proteolysis, thereby rendering them more stable (**Carulli & Verhaagen, 2021; Fawcett & Kwok, 2022; Kitagawa, 2016; Miyata et al., 2012; Miyata & Kitagawa, 2016**). In the zebrafish, expression of 4-O- and 6-O-sulfotransferases are also seen during development (**Chen et al., 2005; Habicher et al., 2015**). Consistent with a potential induced-PNN maturation in early juvenile zebrafish which could be responsible for the behavioral and physiological changes we observed after MMP-2/9 inhibition, is the increase in 4-sulfated CSPGs with no change in 6-sulfated CSPGs suggesting an increase in the 4/6-sulfation ratio known to occur in the transition from juvenile to adults in mammals (**Hou et al., 2017; Miyata et al., 2012**). MMP-2/9 inhibition could also indirectly influence the expression of chst11, a 4-sulfotransferase that may be regulated by transforming growth factor β and arylsulfatase levels. Overall, we hypothesize that MMP-2/9 inhibition increases 4-*O*-sulfated CSPGs aiding to the premature maturation of PNN during the early stages of the juvenile period promoting behavioral stability during learning.

Though our discussion is focused on PNN modulation by inhibition of MMP-2/9, the proteomics data reveal other potential non-mutually exclusive mechanisms by which SB-3CT may affect neuronal excitability and behavior in early juvenile zebrafish. As an example, early juvenile zebrafish treated with SB-3CT exhibit a decrease in neuronal pentraxin receptor b. The pentraxin family is crucial in modulating E/I balance in the hippocampus by modulating PV+ neuronal excitability and the recruitment of pyramidal neurons (**Pelkey et al., 2015**). Indeed, a recent preprint showed that fluoxetine, which in contrast to SB-3CT may increase MMP activity, increased neuronal pentraxin receptor expression in PV+ neurons and also reduced PV expression (**Jetsonen et al., 2024**). It was suggested that reduced PV expression could represent reduced activity of PV neurons. Of additional interest, SB-3CT increased the expression of cathepsin B in late/early adult zebrafish. Though they can target intracellular proteins, cathepsins can also be secreted. Secreted cathepsins can increase MMP-9 activity (**Padamsey et al., 2017**) to modulate CSPGs and PNNs (**Pantazopoulos et al., 2020; Tran et al., 2018**). Increased cathepsin levels may thus represent a compensatory change that occurs after MMP-2/9 inhibition in adult zebrafish. Our proteomic data, though informative, does not provide region specific changes, known to be crucial for circuit plasticity. In this manner, in depth studies parsing out the architecture of the hippocampus and amygdala homologues in zebrafish, for instance, are crucial in determining neuron specific PNN modulations in altering learning and memory and social cognition.

In conclusion, this study highlights the role of MMP-2/9 activity in locomotion, fear memory, sociability, E/I balance, and neuronal population activity in the developing zebrafish brain. It further suggests that MMP-2/9-dependent effects on these endpoints, which have been observed in mammals, are conserved in zebrafish. Moreover, our study demonstrates that as in mammals, zebrafish have CSPG-rich structures surrounding the soma of hippocampal cell bodies that can be modulated by MMPs and/or its downstream substrates. A graphical abstract is provided of hypothetical mechanisms by which MMP-2/9 inhibition affects behavior and neuronal population dynamics in early juvenile zebrafish (Supplementary Fig. S5). Overall, these data posit zebrafish as a useful model to study behavioral and physiological endpoints associated with mental health and neurological disorders as well as adolescent stress, in which proteases and/or their substrates are dysregulated.

## ACKNOWLEDGMENTS

This work was supported by NINDS 5T32NS041218 (IB), F99NS130872 (IB), and R21MH118749 (KC). We would like to acknowledge Dr. Jian-Young Wu for his help with electrophysiology experiments and Dr. Junfeng Ma, the director of the GUMC-Proteomics Shared Resource partially supported by NIH/NCI grant P30 CA051008. The instrument Orbitrap Lumos Tribrid mass spectrometer was partially supported by Dekelbaum Foundation. We apologize to the investigators whose research work may not have been cited.

**No conflict of interest.**

---

**Fig. S1:**
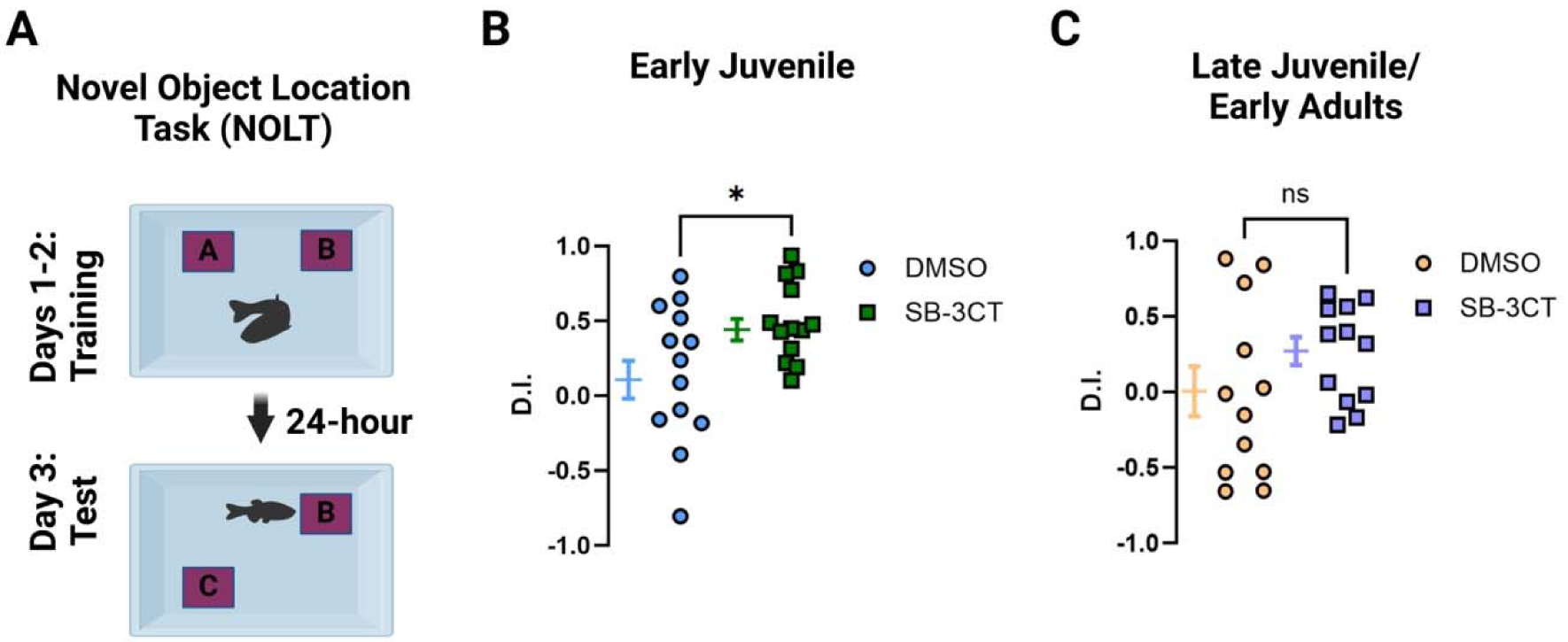
MMP-9 inhibition enhances familiar location recognition in early juvenile zebrafish. **A** Representative cartoon of the Novel Objection Location Task (NOLT) behavioral setup. Similar objects were located on opposing corners (A and B) of the fish tank and time spent in the familiar (corner B) versus novel quadrant (corner C) was counted as exploration time. D.I. formula was defined as time in familiar (F) quadrant minus time in novel (N) quadrant divided by total time in both quadrants. **(B-C)** Show the D.I. of time spent in the familiar quadrant. Graphs are expressed as mean ± S.E.M. Unpaired Student’s t test, *p < 0.05.

**Fig. S2:**
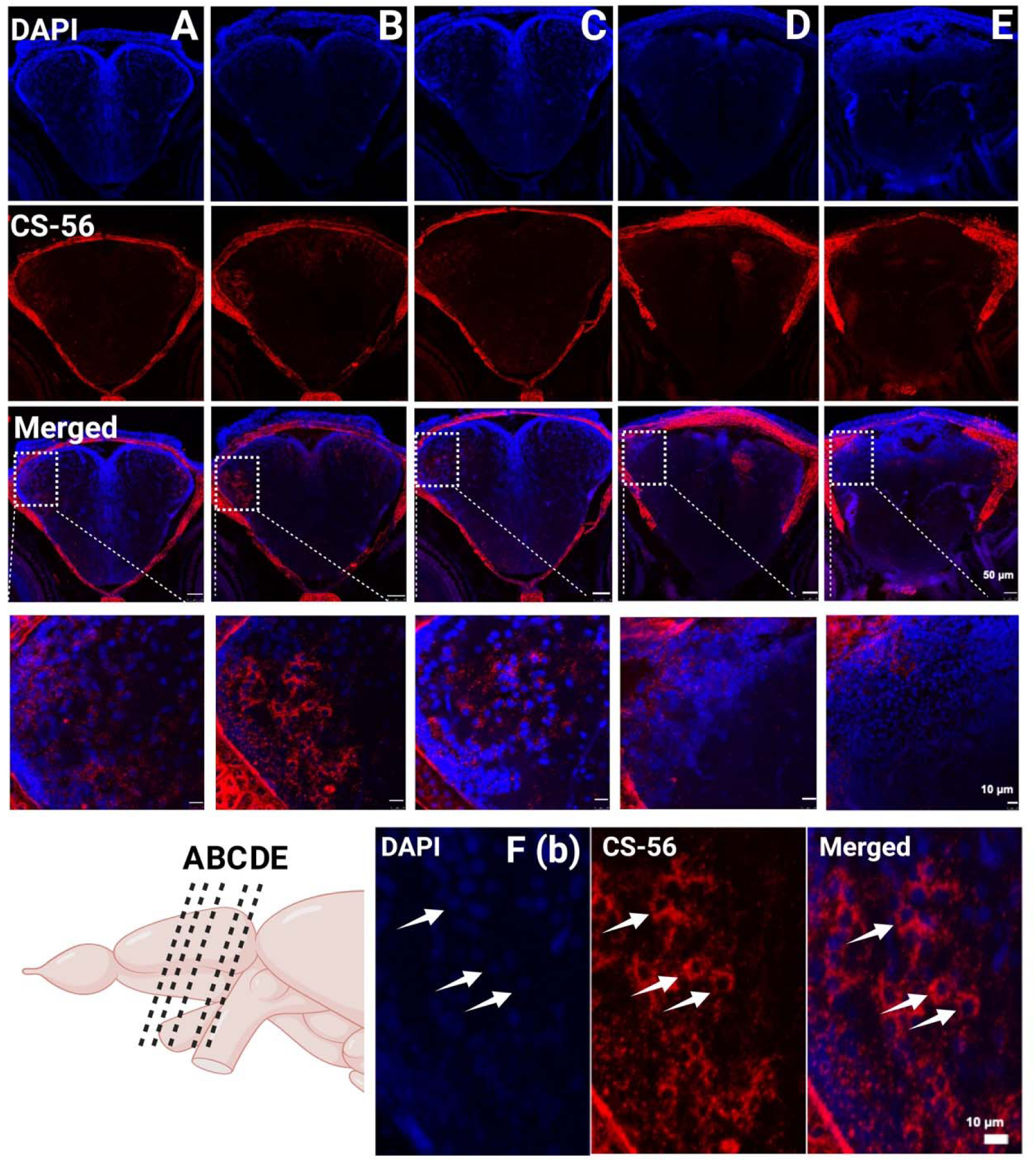
Presence of CS-56+ PNNs in the telencephalon of early juvenile zebrafish. **(A-E)** Show representative images of CS-56+ PNN in the dorsolateral lobe (white squares) of the zebrafish telencephalon. A cartoon showing the lateral view of a zebrafish telencephalon from which the coronal slices were taken (20 μm) is shown in the lower left corner. **F** Shows zoomed-in images from panel B with arrows pointing to cells surrounded by CS-56+ PNNs.

**Fig. S3:**
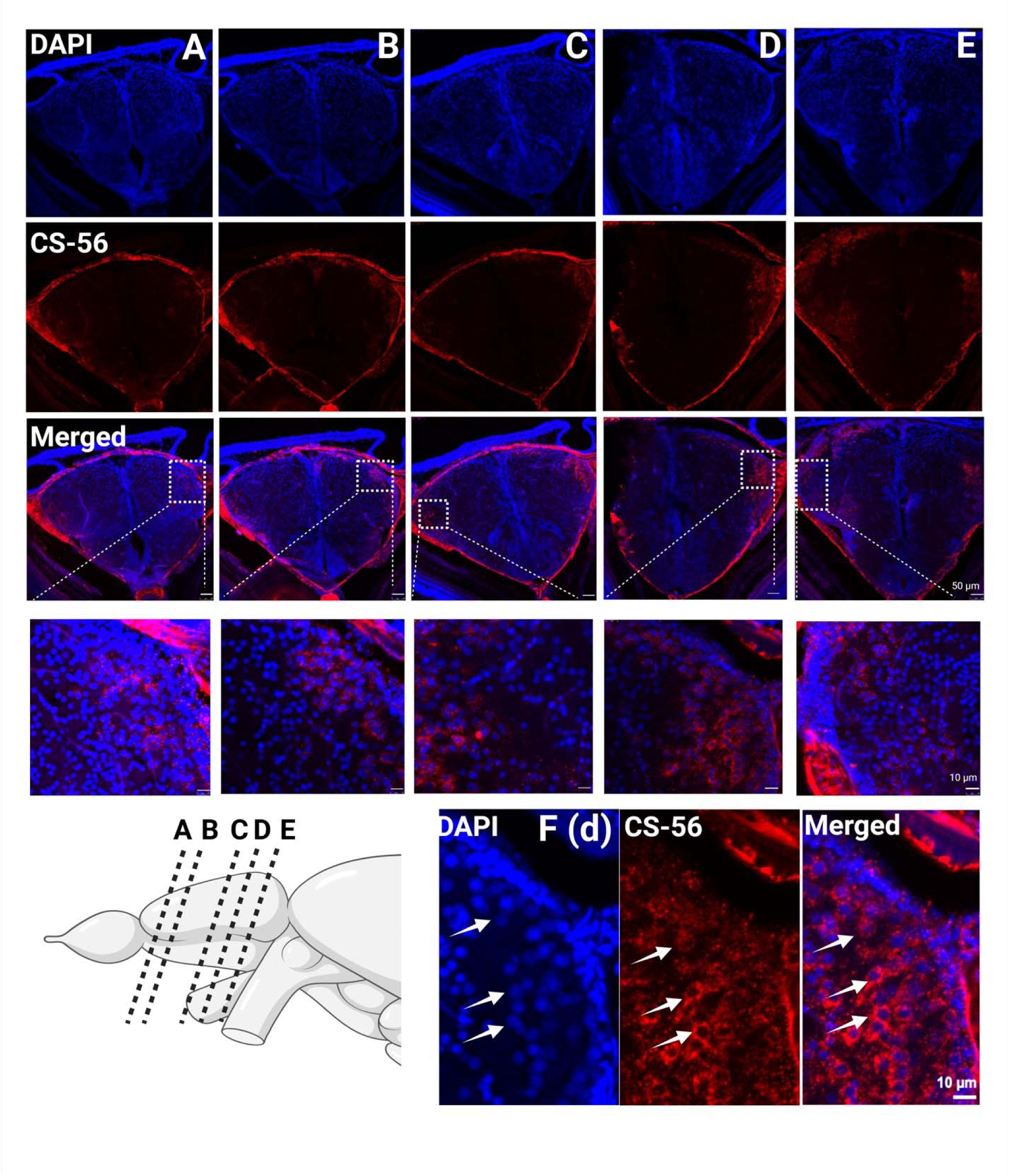
Presence of CS-56+ PNNs in the telencephalon of late juvenile/early adult zebrafish. **(A-E)** Show representative images of CS-56+ PNN in the dorsolateral region of the telencephalon (white squares). A cartoon showing the lateral view of a zebrafish telencephalon from which the coronal slices were taken (20 μm) is shown in the lower left corner. **F** Shows zoomed-in images from panel D with arrows pointing to cells surrounded by CS-56+ PNNs.

**Fig. S4:**
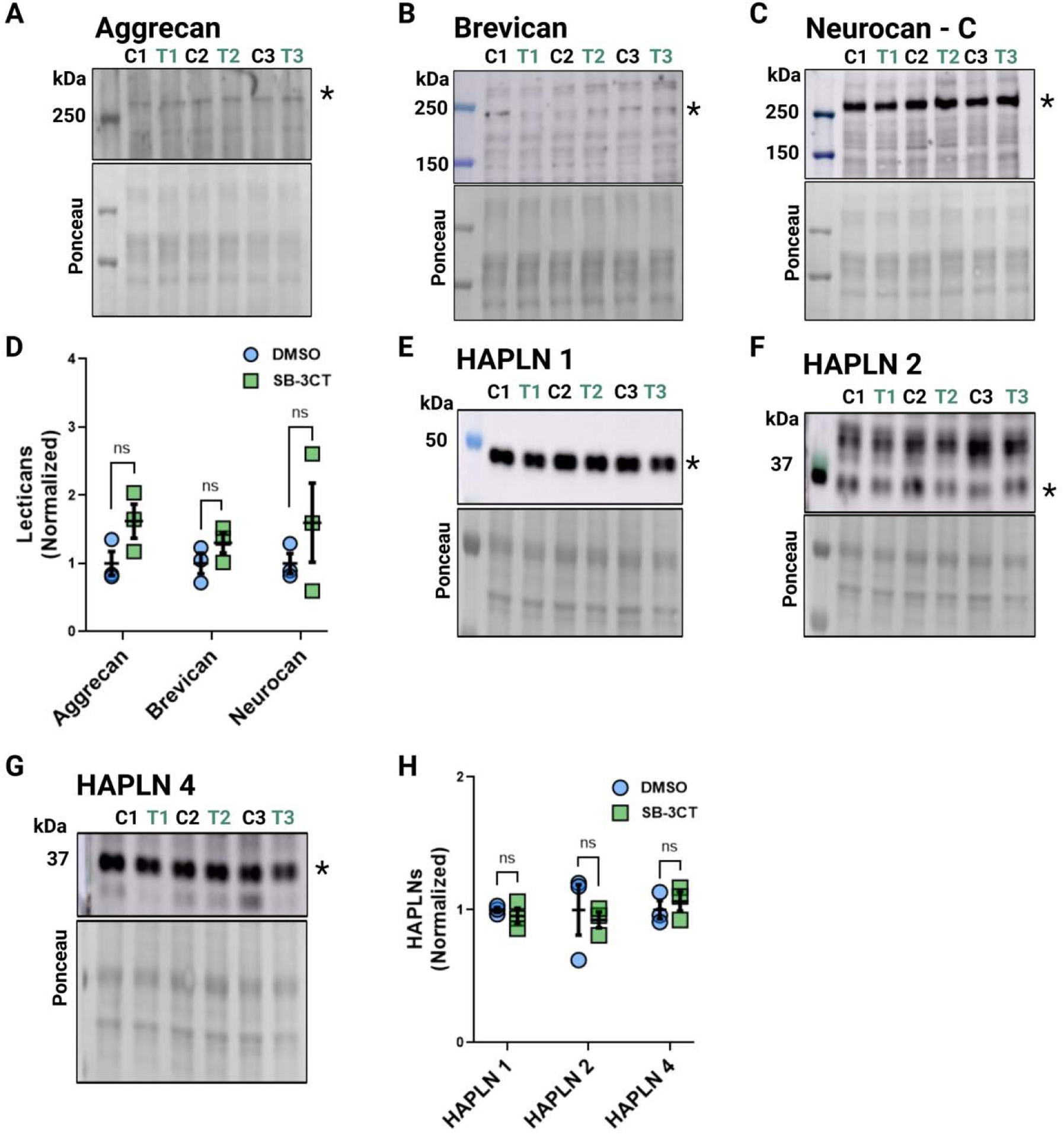
MMP-2/9 inhibition does not alter PNN-core components or HAPLNs in the telencephalon of early juvenile zebrafish. **(A-C)** Representative telencephalon-only pooled Western blots for Aggrecan, Brevican, and Neurocan C, respectively (10-15 telencephalon per group, n=3). Lysates were first digested with ChABC. **D** Quantification of Western blots. **(E-G)** Representative Western blots for 1, 2 and 4 HAPLNs. HAPLN 1 blot was stripped and reprobed for HAPLN 2. **H** Quantification of Western blots. Quantified bands are highlighted using asterisks. Graphs are expressed as mean ± S.E.M. Unpaired t-test. *p < 0.05, ** p < 0.01.

**Fig S5.**
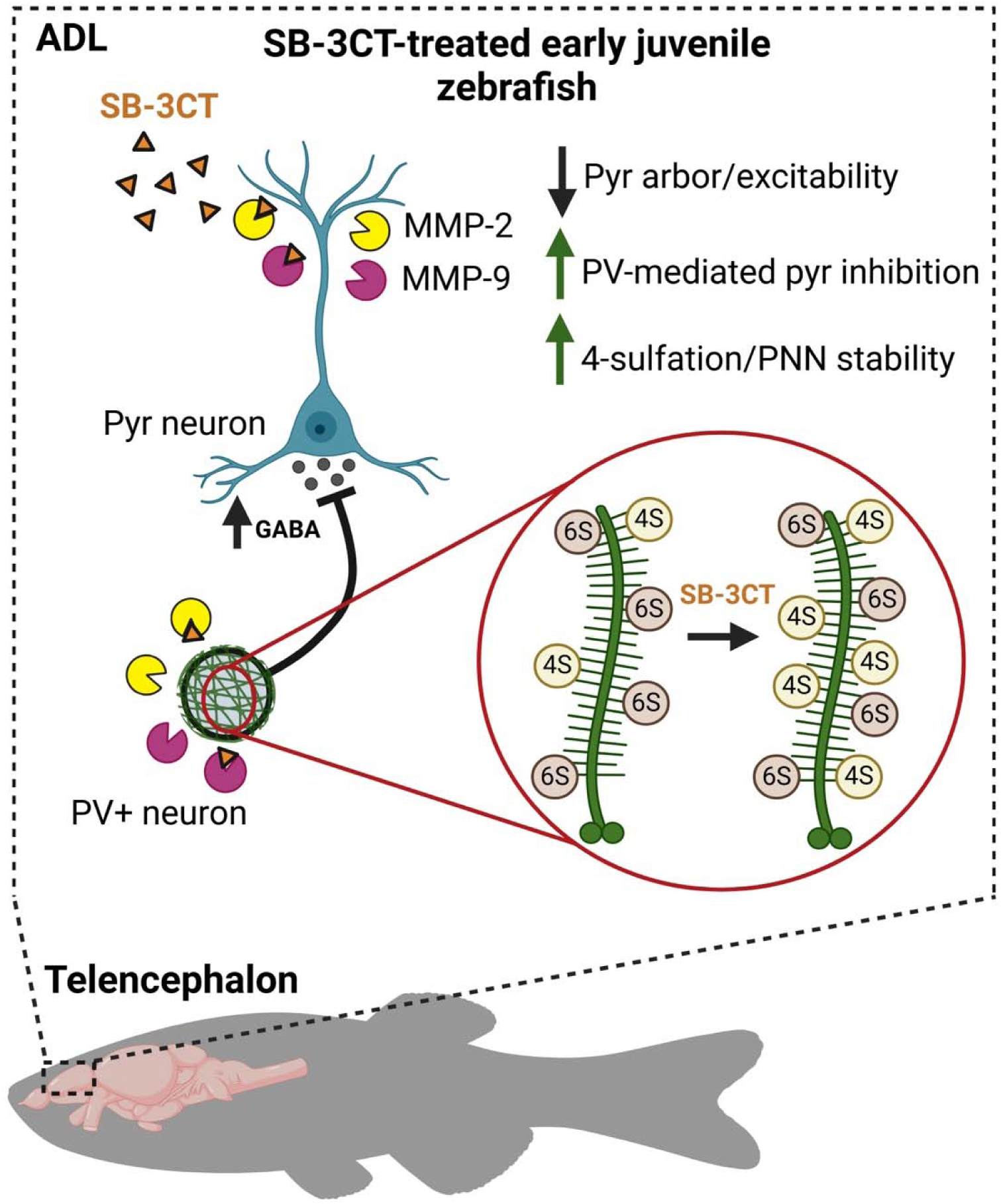
Graphical abstract: SB-3CT decreases MMP-2/9 activity which has been previously shown to decrease dendritic arbor and increase GABAergic-mediated inhibition. The increase in GABAergic inhibition in the ADL of early juvenile zebrafish may be due to a shift in PNN maturation as suggested by an increase in 4-sulfation (4S) known to be increased during adulthood and stabilize these structures against cleavage. Increased PNN expression can enhance fear memory. Together, a reduction in pyramidal (pyr) excitability through enhanced GABAergic transmission may also underline the changes in SWR abundance.

**Table S1:**
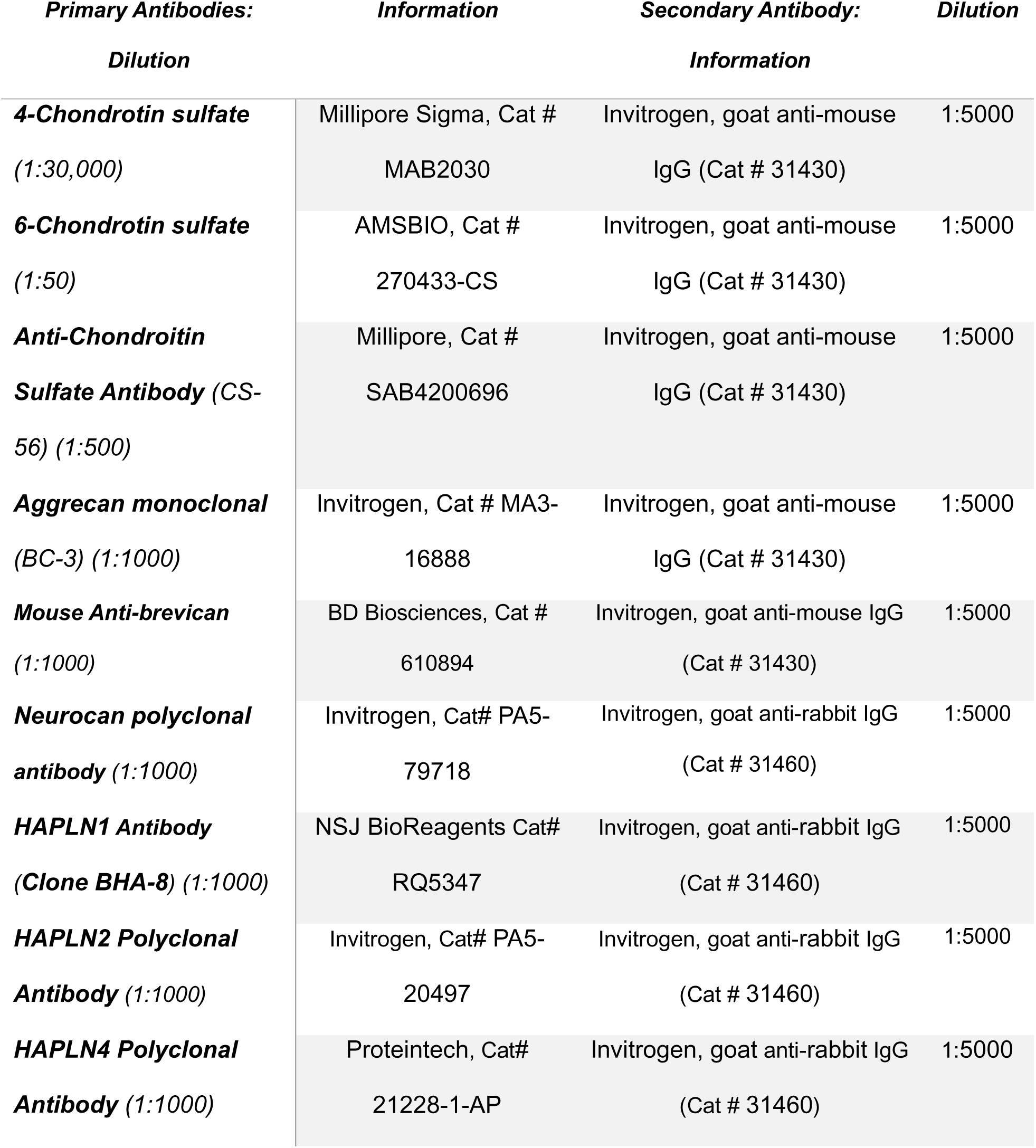
Antibody list.

**Table S2:**
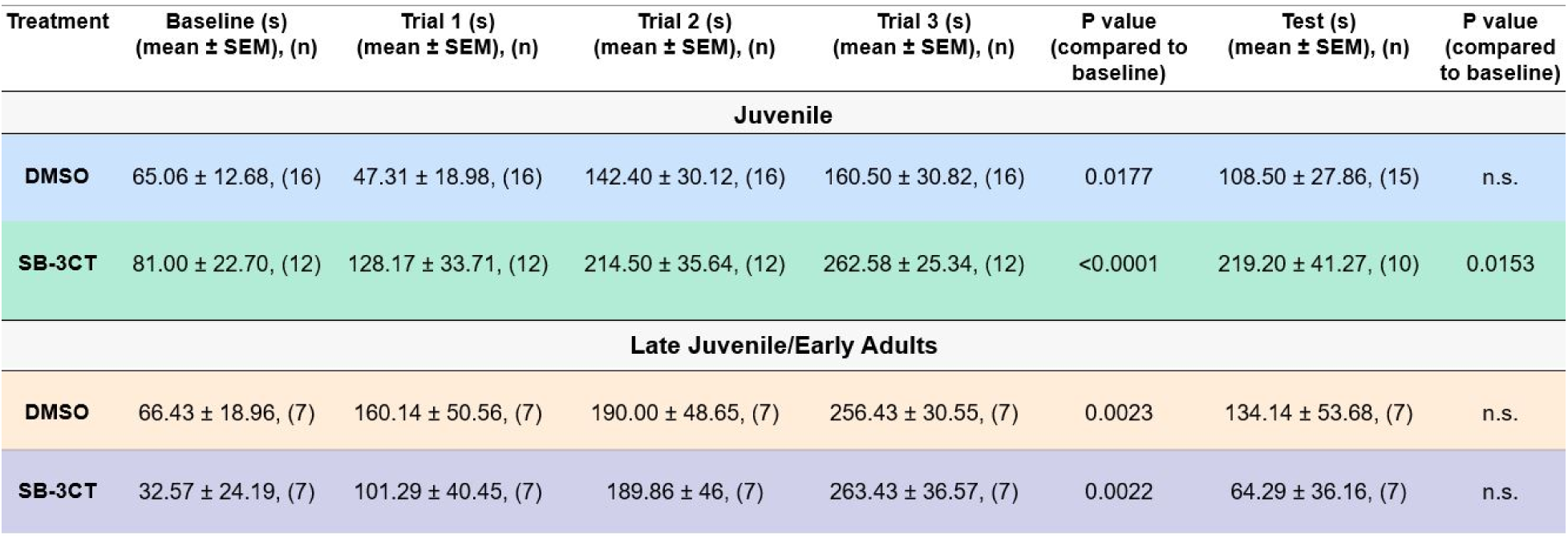
Descriptive statistics for the passive avoidance test (Fig. 2).

| Treatment | Baseline (s)<br>(mean ± SEM), (n) | Trial 1 (s)<br>(mean ± SEM), (n) | Trial 2 (s)<br>(mean ± SEM), (n) | Trial 3 (s)<br>(mean ± SEM), (n) | P value<br>(compared to<br>baseline) | Test (s)<br>(mean ± SEM), (n) | P value<br>(compared<br>to baseline) |
| --- | --- | --- | --- | --- | --- | --- | --- |
| **Juvenile** |  |  |  |  |  |  |  |
| **DMSO** | 65.06 ± 12.68, (16) | 47.31 ± 18.98, (16) | 142.40 ± 30.12, (16) | 160.50 ± 30.82, (16) | 0.0177 | 108.50 ± 27.86, (15) | n.s. |
| **SB-3CT** | 81.00 ± 22.70, (12) | 128.17 ± 33.71, (12) | 214.50 ± 35.64, (12) | 262.58 ± 25.34, (12) | <0.0001 | 219.20 ± 41.27, (10) | 0.0153 |
| **Late Juvenile/Early Adults** |  |  |  |  |  |  |  |
| **DMSO** | 66.43 ± 18.96, (7) | 160.14 ± 50.56, (7) | 190.00 ± 48.65, (7) | 256.43 ± 30.55, (7) | 0.0023 | 134.14 ± 53.68, (7) | n.s. |
| **SB-3CT** | 32.57 ± 24.19, (7) | 101.29 ± 40.45, (7) | 189.86 ± 46, (7) | 263.43 ± 36.57, (7) | 0.0022 | 64.29 ± 36.16, (7) | n.s. |

**Table S3:** Pooled absolute values for each SW duration cluster (Fig. 4).

| Treatment | SW Duration (ms)<br>(mean ± SEM), (n) | P value | Cluster 1 (n, %)<br>(50 – 250 ms) | Cluster 2 (n, %)<br>(300 – 600 ms) | Cluster 3 (n, %)<br>(650 – 1050 ms) | Cluster 4 (n, %)<br>(1075 – 1600 ms) | Total<br>number of<br>events |
| --- | --- | --- | --- | --- | --- | --- | --- |
| Juvenile |  |  |  |  |  |  |  |
| DMSO | 310.9 ± 18.60, (29) | 0.0002 | 14 242, 64.1 | 4 247, 19.1 | 3 193, 14.4 | 548, 2.5 | 22 230 |
| SB3-CT | 429.1 ± 23.16, (29) |  | 7 635, 50.5 | 2 098, 13.9 | 4 642, 30.7<br>2.1-fold increase | 742, 4.9<br>2-fold increase | 15 117 |
| Adults |  |  |  |  |  |  |  |
| DMSO | 266.6 ± 41.52, (12) | 0.8573 | 5 763, 66.1 | 1 605, 18.4 | 1 145, 13.1 | 208, 2.4 | 8 721 |
| SB-3CT | 276.4 ± 33.70, (12) |  | 5 436, 64.9 | 1 740, 20.8 | 988, 11.8 | 218, 2.6 | 8 382 |

## References

Alaiyed, S., Bozzelli, P. L., Caccavano, A., Wu, J. Y., & Conant, K. (2019). Venlafaxine stimulates PNN proteolysis and MMP -9-dependent enhancement of gamma power; relevance to antidepressant efficacy. Journal of Neurochemistry, 148(6), 810–821. 10.1111/jnc.14671

Alaiyed, S., McCann, M., Mahajan, G., Rajkowska, G., Stockmeier, C. A., Kellar, K. J., Wu, J. Y., & Conant, K. (2020). Venlafaxine Stimulates an MMP-9-Dependent Increase in Excitatory/Inhibitory Balance in a Stress Model of Depression. The Journal of Neuroscience, 40(22), 4418–4431. 10.1523/JNEUROSCI.2387-19.2020

Bach, D. R., Tzovara, A., & Vunder, J. (2018). Blocking human fear memory with the matrix metalloproteinase inhibitor doxycycline. Molecular Psychiatry, 23(7), 1584–1589. 10.1038/mp.2017.65

Banerjee, S. B., Gutzeit, V. A., Baman, J., Aoued, H. S., Doshi, N. K., Liu, R. C., & Ressler, K. J. (2017). Perineuronal Nets in the Adult Sensory Cortex Are Necessary for Fear Learning. Neuron, 95(1), 169–179.e3. 10.1016/j.neuron.2017.06.007

Bashirzade, A. A., Zabegalov, K. N., Volgin, A. D., Belova, A. S., Demin, K. A., de Abreu, M. S., Babchenko, V. Ya., Bashirzade, K. A., Yenkoyan, K. B., Tikhonova, M. A., Amstislavskaya, T. G., & Kalueff, A. V. (2022). Modeling neurodegenerative disorders in zebrafish. Neuroscience & Biobehavioral Reviews, 138, 104679. 10.1016/j.neubiorev.2022.104679

Battaglia, F. P., Sutherland, G. R., & McNaughton, B. L. (2004). Hippocampal sharp wave bursts coincide with neocortical “up-state” transitions. Learning & Memory, 11(6), 697–704. 10.1101/lm.73504

Belliveau, C., Théberge, S., Netto, S., Rahimian, R., Fakhfouri, G., Hosdey, C., Davoli, M. A., Hendrickson, A., Hao, K., Giros, B., Turecki, G., Alonge, K. M., & Mechawar, N. (2024). Chondroitin sulfate glycan sulfation patterns influence histochemical labeling of perineuronal nets: A comparative study of interregional distribution in human and mouse brain. Glycobiology, 34(8), cwae049. 10.1093/glycob/cwae049

Blanco, I., Caccavano, A., Wu, J.-Y., Vicini, S., Glasgow, E., & Conant, K. (2024). Coupling of Sharp Wave Events between Zebrafish Hippocampal and Amygdala Homologues. The Journal of Neuroscience, e1467232024. 10.1523/JNEUROSCI.1467-23.2024

Born, J. (2010). Slow-wave sleep and the consolidation of long-term memory. The World Journal of Biological Psychiatry, 11(sup1), 16–21. 10.3109/15622971003637637

Brenet, A., Hassan-Abdi, R., Somkhit, J., Yanicostas, C., & Soussi-Yanicostas, N. (2019). Defective Excitatory/Inhibitory Synaptic Balance and Increased Neuron Apoptosis in a Zebrafish Model of Dravet Syndrome. Cells, 8(10), 1199. 10.3390/cells8101199

Burmeister, M., Fraunenstein, A., Kahms, M., Arends, L., Gerwien, H., Deshpande, T., Kuhlmann, T., Gross, C. C., Naik, V. N., Wiendl, H., Klingauf, J., Meissner, F., & Sorokin, L. (2023). Secretomics reveals gelatinase substrates at the blood-brain barrier that are implicated in astroglial barrier function. Science Advances, 9(29), eadg0686. 10.1126/sciadv.adg0686

Buzsáki, G. (2015). Hippocampal sharp wave-ripple: A cognitive biomarker for episodic memory and planning. Hippocampus, 25(10), 1073–1188. 10.1002/hipo.22488

Cabungcal, J.-H., Steullet, P., Morishita, H., Kraftsik, R., Cuenod, M., Hensch, T. K., & Do, K. Q. (2013). Perineuronal nets protect fast-spiking interneurons against oxidative stress. Proceedings of the National Academy of Sciences, 110(22), 9130–9135. 10.1073/pnas.1300454110

Caccavano, A., Bozzelli, P. L., Forcelli, P. A., Pak, D. T. S., Wu, J.-Y., Conant, K., & Vicini, S. (2020). Inhibitory Parvalbumin Basket Cell Activity is Selectively Reduced during Hippocampal Sharp Wave Ripples in a Mouse Model of Familial Alzheimer’s Disease. The Journal of Neuroscience, 40(26), 5116–5136. 10.1523/JNEUROSCI.0425-20.2020

Carstens, K. E., Lustberg, D. J., Shaughnessy, E. K., McCann, K. E., Alexander, G. M., & Dudek, S. M. (2021). Perineuronal net degradation rescues CA2 plasticity in a mouse model of Rett syndrome. Journal of Clinical Investigation, 131(16), e137221. 10.1172/JCI137221

Carstens, K. E., Phillips, M. L., Pozzo-Miller, L., Weinberg, R. J., & Dudek, S. M. (2016). Perineuronal Nets Suppress Plasticity of Excitatory Synapses on CA2 Pyramidal Neurons. The Journal of Neuroscience, 36(23), 6312–6320. 10.1523/JNEUROSCI.0245-16.2016

Carulli, D., Pizzorusso, T., Kwok, J. C. F., Putignano, E., Poli, A., Forostyak, S., Andrews, M. R., Deepa, S. S., Glant, T. T., & Fawcett, J. W. (2010). Animals lacking link protein have attenuated perineuronal nets and persistent plasticity. Brain, 133(8), 2331–2347. 10.1093/brain/awq145

Carulli, D., & Verhaagen, J. (2021). An Extracellular Perspective on CNS Maturation: Perineuronal Nets and the Control of Plasticity. International Journal of Molecular Sciences, 22(5), 2434. 10.3390/ijms22052434

Chen, E., Stringer, S. E., Rusch, M. A., Selleck, S. B., & Ekker, S. C. (2005). A unique role for 6-O sulfation modification in zebrafish vascular development. Developmental Biology, 284(2), 364–376. 10.1016/j.ydbio.2005.05.032

Conant, K., Allen, M., & Lim, S. T. (2015). Activity dependent CAM cleavage and neurotransmission. Frontiers in Cellular Neuroscience, 9. 10.3389/fncel.2015.00305

Cope, E. C., Zych, A. D., Katchur, N. J., Waters, R. C., Laham, B. J., Diethorn, E. J., Park, C. Y., Meara, W. R., & Gould, E. (2022). Atypical perineuronal nets in the CA2 region interfere with social memory in a mouse model of social dysfunction. Molecular Psychiatry, 27(8), 3520–3531. 10.1038/s41380-021-01174-2

Coutellier, L., & Usdin, T. B. (2011). Enhanced long-term fear memory and increased anxiety and depression-like behavior after exposure to an aversive event in mice lacking TIP39 signaling. Behavioural Brain Research, 222(1), 265–269. 10.1016/j.bbr.2011.02.043

Davidson, T. J., Kloosterman, F., & Wilson, M. A. (2009). Hippocampal Replay of Extended Experience. Neuron, 63(4), 497–507. 10.1016/j.neuron.2009.07.027

Domínguez, S., Rey, C. C., Therreau, L., Fanton, A., Massotte, D., Verret, L., Piskorowski, R. A., & Chevaleyre, V. (2019). Maturation of PNN and ErbB4 Signaling in Area CA2 during Adolescence Underlies the Emergence of PV Interneuron Plasticity and Social Memory. Cell Reports, 29(5), 1099–1112.e4. 10.1016/j.celrep.2019.09.044

Fawcett, J. W., Fyhn, M., Jendelova, P., Kwok, J. C. F., Ruzicka, J., & Sorg, B. A. (2022). The extracellular matrix and perineuronal nets in memory. Molecular Psychiatry, 27(8), 3192–3203. 10.1038/s41380-022-01634-3

Fawcett, J. W., & Kwok, J. C. F. (2022). Proteoglycan Sulphation in the Function of the Mature Central Nervous System. Frontiers in Integrative Neuroscience, 16, 895493. 10.3389/fnint.2022.895493

Fernández-Ruiz, A., Oliva, A., Fermino de Oliveira, E., Rocha-Almeida, F., Tingley, D., & Buzsáki, G. (2019). Long-duration hippocampal sharp wave ripples improve memory. Science, 364(6445), 1082–1086. 10.1126/science.aax0758

Folgueira, M., Bayley, P., Navratilova, P., Becker, T. S., Wilson, S. W., & Clarke, J. D. (2012). Morphogenesis underlying the development of the everted teleost telencephalon. Neural Development, 7(1), 212. 10.1186/1749-8104-7-32

Frischknecht, R., Heine, M., Perrais, D., Seidenbecher, C. I., Choquet, D., & Gundelfinger, E. D. (2009). Brain extracellular matrix affects AMPA receptor lateral mobility and short-term synaptic plasticity. Nature Neuroscience, 12(7), 897–904. 10.1038/nn.2338

Ganz, J., Kroehne, V., Freudenreich, D., Machate, A., Geffarth, M., Braasch, I., Kaslin, J., & Brand, M. (2015). Subdivisions of the adult zebrafish pallium based on molecular marker analysis. F1000Research, 3, 308. 10.12688/f1000research.5595.2

Giri, B., Kinsky, N., Kaya, U., Maboudi, K., Abel, T., & Diba, K. (2024). Sleep loss diminishes hippocampal reactivation and replay. Nature, 630(8018), 935–942. 10.1038/s41586-024-07538-2

Gogolla, N., Caroni, P., Lüthi, A., & Herry, C. (2009). Perineuronal Nets Protect Fear Memories from Erasure. Science, 325(5945), 1258–1261. 10.1126/science.1174146

Gorkiewicz, T., Balcerzyk, M., Kaczmarek, L., & Knapska, E. (2015). Matrix metalloproteinase 9 (MMP-9) is indispensable for long term potentiation in the central and basal but not in the lateral nucleus of the amygdala. Frontiers in Cellular Neuroscience, 9. 10.3389/fncel.2015.00073

Grochecki, P., Smaga, I., Wydra, K., Marszalek-Grabska, M., Slowik, T., Kedzierska, E., Listos, J., Gibula-Tarlowska, E., Filip, M., & Kotlinska, J. H. (2023). Impact of Mephedrone on Fear Memory in Adolescent Rats: Involvement of Matrix Metalloproteinase-9 (MMP-9) and N-Methyl-D-aspartate (NMDA) Receptor. International Journal of Molecular Sciences, 24(3), 1941. 10.3390/ijms24031941

Habicher, J., Filipek-Górniok, B., Kjellén, L., & Ledin, J. (2021). Proteoglycans in Zebrafish Development. In M. Götte & K. Forsberg-Nilsson (Eds.), Proteoglycans in Stem Cells (Vol. 9, pp. 21–34). Springer International Publishing. 10.1007/978-3-030-73453-4_2

Habicher, J., Haitina, T., Eriksson, I., Holmborn, K., Dierker, T., Ahlberg, P. E., & Ledin, J. (2015). Chondroitin / Dermatan Sulfate Modification Enzymes in Zebrafish Development. PLOS ONE, 10(3), e0121957. 10.1371/journal.pone.0121957

Happel, M. F. K., Niekisch, H., Castiblanco Rivera, L. L., Ohl, F. W., Deliano, M., & Frischknecht, R. (2014). Enhanced cognitive flexibility in reversal learning induced by removal of the extracellular matrix in auditory cortex. Proceedings of the National Academy of Sciences, 111(7), 2800–2805. 10.1073/pnas.1310272111

Härtig, W., Meinicke, A., Michalski, D., Schob, S., & Jäger, C. (2022). Update on Perineuronal Net Staining With Wisteria floribunda Agglutinin (WFA). Frontiers in Integrative Neuroscience, 16, 851988. 10.3389/fnint.2022.851988

Hernandez-Anzaldo, S., Brglez, V., Hemmeryckx, B., Leung, D., Filep, J. G., Vance, J. E., Vance, D. E., Kassiri, Z., Lijnen, R. H., Lambeau, G., & Fernandez-Patron, C. (2016). Novel Role for Matrix Metalloproteinase 9 in Modulation of Cholesterol Metabolism. Journal of the American Heart Association, 5(10). 10.1161/JAHA.116.004228

Hou, X., Yoshioka, N., Tsukano, H., Sakai, A., Miyata, S., Watanabe, Y., Yanagawa, Y., Sakimura, K., Takeuchi, K., Kitagawa, H., Hensch, T. K., Shibuki, K., Igarashi, M., & Sugiyama, S. (2017). Chondroitin Sulfate Is Required for Onset and Offset of Critical Period Plasticity in Visual Cortex. Scientific Reports, 7(1), 12646. 10.1038/s41598-017-04007-x

Huang, H., Joffrin, A. M., Zhao, Y., Miller, G. M., Zhang, G. C., Oka, Y., & Hsieh-Wilson, L. C. (2023). Chondroitin 4-*O* -sulfation regulates hippocampal perineuronal nets and social memory. Proceedings of the National Academy of Sciences, 120(24), e2301312120. 10.1073/pnas.2301312120

Hylin, M. J., Orsi, S. A., Moore, A. N., & Dash, P. K. (2013). Disruption of the perineuronal net in the hippocampus or medial prefrontal cortex impairs fear conditioning. Learning & Memory, 20(5), 267–273. 10.1101/lm.030197.112

Irvine, S., & Kwok, J. (2018). Perineuronal Nets in Spinal Motoneurones: Chondroitin Sulphate Proteoglycan around Alpha Motoneurones. International Journal of Molecular Sciences, 19(4), 1172. 10.3390/ijms19041172

Jetsonen, E., Suleymanova, I., Castrén, E., & Umemori, J. (2024). Chronic treatment with fluoxetine downregulates mitochondrial activity in parvalbumin interneurons of prefrontal cortex. bioRxiv. 10.1101/2024.09.27.615344

Jovasevic, V., Zhang, H., Petrovic, Z., Cicvaric, A., & Radulovic, J. (2021). Protocol for assessing the role of hippocampal perineuronal nets in aversive memories. STAR Protocols, 2(4), 100931. 10.1016/j.xpro.2021.100931

Kalueff, A. V., Stewart, A. M., & Gerlai, R. (2014). Zebrafish as an emerging model for studying complex brain disorders. Trends in Pharmacological Sciences, 35(2), 63–75. 10.1016/j.tips.2013.12.002

Kennedy-Wood, K., Ng, C. A. S., Alaiyed, S., Foley, P. L., & Conant, K. (2021). Increased MMP-9 levels with strain-dependent stress resilience and tunnel handling in mice. Behavioural Brain Research, 408, 113288. 10.1016/j.bbr.2021.113288

Kim, Y.-H., Lee, Y., Kim, D., Jung, M. W., & Lee, C.-J. (2010). Scopolamine-induced learning impairment reversed by physostigmine in zebrafish. Neuroscience Research, 67(2), 156–161. 10.1016/j.neures.2010.03.003

Kitagawa, H. (2016). Chondroitin sulfate and neuronal disorders. Frontiers in Bioscience, 21(7), 1330–1340. 10.2741/4460

Knapska, E., & Kaczmarek, L. (2016). Matrix Metalloproteinase 9 (MMP-9) in Learning and Memory. In K. P. Giese & K. Radwanska (Eds.), Novel Mechanisms of Memory (pp. 161–181). Springer International Publishing. 10.1007/978-3-319-24364-1_9

Koyuncu, A., İnce, E., Ertekin, E., & Tükel, R. (2019). Comorbidity in social anxiety disorder: Diagnostic and therapeutic challenges. Drugs in Context, 8, 1–13. 10.7573/dic.212573

LeBert, D. C., Squirrell, J. M., Rindy, J., Broadbridge, E., Lui, Y., Zakrzewska, A., Eliceiri, K. W., Meijer, A. H., & Huttenlocher, A. (2015). Matrix metalloproteinase 9 modulates collagen matrices and wound repair. Development, 142(12), 2136–2146. 10.1242/dev.121160

Lee, S.-J., Wei, M., Zhang, C., Maxeiner, S., Pak, C., Calado Botelho, S., Trotter, J., Sterky, F. H., & Südhof, T. C. (2017). Presynaptic Neuronal Pentraxin Receptor Organizes Excitatory and Inhibitory Synapses. The Journal of Neuroscience, 37(5), 1062–1080. 10.1523/JNEUROSCI.2768-16.2016

Mattioni, L., Barbieri, A., Grigoli, A., Balasco, L., Bozzi, Y., & Provenzano, G. (2024). Alterations of Perineuronal Net Expression and Abnormal Social Behavior and Whisker-dependent Texture Discrimination in Mice Lacking the Autism Candidate Gene Engrailed 2. Neuroscience, 546, 63–74. 10.1016/j.neuroscience.2024.03.023

Meisel, J. E., & Chang, M. (2017). Selective small-molecule inhibitors as chemical tools to define the roles of matrix metalloproteinases in disease. Biochimica et Biophysica Acta (BBA) - Molecular Cell Research, 1864(11), 2001–2014. 10.1016/j.bbamcr.2017.04.011

Meyer-Puttlitz, B., Milev, P., Junker, E., Zimmer, I., Margolis, R. U., & Margolis, R. K. (2002). Chondroitin Sulfate and Chondroitin/Keratan Sulfate Proteoglycans of Nervous Tissue: Developmental Changes of Neurocan and Phosphacan. Journal of Neurochemistry, 65(5), 2327–2337. 10.1046/j.1471-4159.1995.65052327.x

Michaluk, P., Kolodziej, L., Mioduszewska, B., Wilczynski, G. M., Dzwonek, J., Jaworski, J., Gorecki, D. C., Ottersen, O. P., & Kaczmarek, L. (2007). β-Dystroglycan as a Target for MMP-9, in Response to Enhanced Neuronal Activity. Journal of Biological Chemistry, 282(22), 16036–16041. 10.1074/jbc.M700641200

Miyata, S., & Kitagawa, H. (2016). Chondroitin 6-Sulfation Regulates Perineuronal Net Formation by Controlling the Stability of Aggrecan. Neural Plasticity, 2016, 1–13. 10.1155/2016/1305801

Miyata, S., Komatsu, Y., Yoshimura, Y., Taya, C., & Kitagawa, H. (2012). Persistent cortical plasticity by upregulation of chondroitin 6-sulfation. Nature Neuroscience, 15(3), 414–422. 10.1038/nn.3023

Mondal, S., Adhikari, N., Banerjee, S., Amin, S. A., & Jha, T. (2020). Matrix metalloproteinase-9 (MMP-9) and its inhibitors in cancer: A minireview. European Journal of Medicinal Chemistry, 194, 112260. 10.1016/j.ejmech.2020.112260

Morikawa, S., Ikegaya, Y., Narita, M., & Tamura, H. (2017). Activation of perineuronal net-expressing excitatory neurons during associative memory encoding and retrieval. Scientific Reports, 7(1), 46024. 10.1038/srep46024

Mu, Y., Bennett, D. V., Rubinov, M., Narayan, S., Yang, C.-T., Tanimoto, M., Mensh, B. D., Looger, L. L., & Ahrens, M. B. (2019). Glia Accumulate Evidence that Actions Are Futile and Suppress Unsuccessful Behavior. Cell, 178(1), 27–43.e19. 10.1016/j.cell.2019.05.050

Nagy, V. (2006). Matrix Metalloproteinase-9 Is Required for Hippocampal Late-Phase Long-Term Potentiation and Memory. Journal of Neuroscience, 26(7), 1923–1934. 10.1523/JNEUROSCI.4359-05.2006

Nguyen, M., Stewart, A. M., & Kalueff, A. V. (2014). Aquatic blues: Modeling depression and antidepressant action in zebrafish. Progress in Neuro-Psychopharmacology and Biological Psychiatry, 55, 26–39. 10.1016/j.pnpbp.2014.03.003

Padamsey, Z., McGuinness, L., Bardo, S. J., Reinhart, M., Tong, R., Hedegaard, A., Hart, M. L., & Emptage, N. J. (2017). Activity-Dependent Exocytosis of Lysosomes Regulates the Structural Plasticity of Dendritic Spines. Neuron, 93(1), 132–146. 10.1016/j.neuron.2016.11.013

Pantazopoulos, H., Gisabella, B., Rexrode, L., Benefield, D., Yildiz, E., Seltzer, P., Valeri, J., Chelini, G., Reich, A., Ardelt, M., & Berretta, S. (2020). Circadian Rhythms of Perineuronal Net Composition. Eneuro, 7(4), ENEURO.0034-19.2020. 10.1523/ENEURO.0034-19.2020

Pelkey, K. A., Barksdale, E., Craig, M. T., Yuan, X., Sukumaran, M., Vargish, G. A., Mitchell, R. M., Wyeth, M. S., Petralia, R. S., Chittajallu, R., Karlsson, R.-M., Cameron, H. A., Murata, Y., Colonnese, M. T., Worley, P. F., & McBain, C. J. (2015). Pentraxins Coordinate Excitatory Synapse Maturation and Circuit Integration of Parvalbumin Interneurons. Neuron, 85(6), 1257–1272. 10.1016/j.neuron.2015.02.020

Pirbhoy, P. S., Rais, M., Lovelace, J. W., Woodard, W., Razak, K. A., Binder, D. K., & Ethell, I. M. (2020). Acute pharmacological inhibition of matrix metalloproteinase-9 activity during development restores perineuronal net formation and normalizes auditory processing in *Fmr1* KO mice. Journal of Neurochemistry, 155(5), 538–558. 10.1111/jnc.15037

Poli, A., Viglione, A., Mazziotti, R., Totaro, V., Morea, S., Melani, R., Silingardi, D., Putignano, E., Berardi, N., & Pizzorusso, T. (2023). Selective Disruption of Perineuronal Nets in Mice Lacking Crtl1 is Sufficient to Make Fear Memories Susceptible to Erasure. Molecular Neurobiology, 60(7), 4105–4119. 10.1007/s12035-023-03314-x

Ramsaran, A. I., Wang, Y., Golbabaei, A., Yeung, B. A., de Snoo, M. L., Rashid, A. J., Awasthi, A., Lau, J., Tran, L. M., Ko, S. Y., Abegg, A., Duan, L. C., McKenzie, C., Gallucci, J., Ahmed, M., Kaushik, R., Dityatev, A., Josselyn, S. A., & Frankland, P. W. (2023). A shift in the mechanisms controlling hippocampal engram formation during brain maturation [Preprint]. Neuroscience. 10.1101/2023.01.09.523283

Reinhard, S. M., Razak, K., & Ethell, I. M. (2015). A delicate balance: Role of MMP-9 in brain development and pathophysiology of neurodevelopmental disorders. Frontiers in Cellular Neuroscience, 9. 10.3389/fncel.2015.00280

Rey, C. C., Robert, V., Bouisset, G., Loisy, M., Lopez, S., Cattaud, V., Lejards, C., Piskorowski, R. A., Rampon, C., Chevaleyre, V., & Verret, L. (2022). Altered inhibitory function in hippocampal CA2 contributes in social memory deficits in Alzheimer’s mouse model. iScience, 25(3), 103895. 10.1016/j.isci.2022.103895

Riga, D., Kramvis, I., Koskinen, M. K., van Bokhoven, P., van der Harst, J. E., Heistek, T. S., Jaap Timmerman, A., van Nierop, P., van der Schors, R. C., Pieneman, A. W., de Weger, A., van Mourik, Y., Schoffelmeer, A. N. M., Mansvelder, H. D., Meredith, R. M., Hoogendijk, W. J. G., Smit, A. B., & Spijker, S. (2017). Hippocampal extracellular matrix alterations contribute to cognitive impairment associated with a chronic depressive-like state in rats. Science Translational Medicine, 9(421), eaai8753. 10.1126/scitranslmed.aai8753

Ruoslahti, E. (1996). Brain extracellular matrix. Glycobiology, 6(5), 489–492. 10.1093/glycob/6.5.489

Ruzicka, J., Dalecka, M., Safrankova, K., Peretti, D., Jendelova, P., Kwok, J. C. F., & Fawcett, J. W. (2022). Perineuronal nets affect memory and learning after synapse withdrawal. Translational Psychiatry, 12(1), 480. 10.1038/s41398-022-02226-z

S., A., P. L., B., A., C., J.Y., W., & K., C. (2018). Venlafaxine stimulates PNN proteolysis and MMP-9 dependent enhancement of gamma power; relevance to antidepressant efficacy [Preprint]. Neuroscience. 10.1101/432419

Sanchez, B., Kraszewski, P., Lee, S., & Cope, E. C. (2023). From molecules to behavior: Implications for perineuronal net remodeling in learning and memory. Journal of Neurochemistry, jnc.16036. 10.1111/jnc.16036

Schuch, F., Vancampfort, D., Firth, J., Rosenbaum, S., Ward, P., Reichert, T., Bagatini, N. C., Bgeginski, R., & Stubbs, B. (2017). Physical activity and sedentary behavior in people with major depressive disorder: A systematic review and meta-analysis. Journal of Affective Disorders, 210, 139–150. 10.1016/j.jad.2016.10.050

Shein-Idelson, M., Ondracek, J. M., Liaw, H.-P., Reiter, S., & Laurent, G. (2016). Slow waves, sharp waves, ripples, and REM in sleeping dragons. Science (New York, N.Y.), 352(6285), 590–595. 10.1126/science.aaf3621

Silva, N. J., Nagashima, M., Li, J., Kakuk-Atkins, L., Ashrafzadeh, M., Hyde, D. R., & Hitchcock, P. F. (2019). Inflammation and matrix metalloproteinase 9 (Mmp-9) regulate photoreceptor regeneration in adult zebrafish [Preprint]. Neuroscience. 10.1101/518365

Stange, J. P., Connolly, S. L., Burke, T. A., Hamilton, J. L., Hamlat, E. J., Abramson, L. Y., & Alloy, L. B. (2016). INFLEXIBLE COGNITION PREDICTS FIRST ONSET OF MAJOR DEPRESSIVE EPISODES IN ADOLESCENCE. Depression and Anxiety, 33(11), 1005–1012. 10.1002/da.22513

Sun, Z. Y., Bozzelli, P. L., Caccavano, A., Allen, M., Balmuth, J., Vicini, S., Wu, J.-Y., & Conant, K. (2018). Disruption of perineuronal nets increases the frequency of sharp wave ripple events. Hippocampus, 28(1), 42–52. 10.1002/hipo.22804

Szklarczyk, A., Lapinska, J., Rylski, M., McKay, R. D. G., & Kaczmarek, L. (2002). Matrix Metalloproteinase-9 Undergoes Expression and Activation during Dendritic Remodeling in Adult Hippocampus. The Journal of Neuroscience, 22(3), 920–930. 10.1523/JNEUROSCI.22-03-00920.2002

Takeda, A., Shuto, M., & Funakoshi, K. (2018). Chondroitin Sulfate Expression in Perineuronal Nets After Goldfish Spinal Cord Lesion. Frontiers in Cellular Neuroscience, 12, 63. 10.3389/fncel.2018.00063

Takiguchi, M., Morinobu, S., & Funakoshi, K. (2021). Chondroitin sulfate expression around spinal motoneurons during postnatal development in rats. Brain Research, 147252. 10.1016/j.brainres.2020.147252

Tewari, B. P., Chaunsali, L., Campbell, S. L., Patel, D. C., Goode, A. E., & Sontheimer, H. (2018). Perineuronal nets decrease membrane capacitance of peritumoral fast spiking interneurons in a model of epilepsy. Nature Communications, 9(1), 4724. 10.1038/s41467-018-07113-0

Tran, A. P., Sundar, S., Yu, M., Lang, B. T., & Silver, J. (2018). Modulation of Receptor Protein Tyrosine Phosphatase Sigma Increases Chondroitin Sulfate Proteoglycan Degradation through Cathepsin B Secretion to Enhance Axon Outgrowth. The Journal of Neuroscience, 38(23), 5399–5414. 10.1523/JNEUROSCI.3214-17.2018

Vargas, R., Þorsteinsson, H., & Karlsson, K. Æ. (2012). Spontaneous neural activity of the anterodorsal lobe and entopeduncular nucleus in adult zebrafish: A putative homologue of hippocampal sharp waves. Behavioural Brain Research, 229(1), 10–20. 10.1016/j.bbr.2011.12.025

Venturino, A., Schulz, R., De Jesús-Cortés, H., Maes, M. E., Nagy, B., Reilly-Andújar, F., Colombo, G., Cubero, R. J. A., Schoot Uiterkamp, F. E., Bear, M. F., & Siegert, S. (2021). Microglia enable mature perineuronal nets disassembly upon anesthetic ketamine exposure or 60-Hz light entrainment in the healthy brain. Cell Reports, 36(1), 109313. 10.1016/j.celrep.2021.109313

Wang, X. -b., Bozdagi, O., Nikitczuk, J. S., Zhai, Z. W., Zhou, Q., & Huntley, G. W. (2008). Extracellular proteolysis by matrix metalloproteinase-9 drives dendritic spine enlargement and long-term potentiation coordinately. Proceedings of the National Academy of Sciences, 105(49), 19520–19525. 10.1073/pnas.0807248105

Wright, J. W., & Harding, J. W. (2009). Contributions of Matrix Metalloproteinases to Neural Plasticity, Habituation, Associative Learning and Drug Addiction. Neural Plasticity, 2009, 1–12. 10.1155/2009/579382

Wyatt, R. A., & Crawford, B. D. (2021). Post-translational activation of Mmp2 correlates with patterns of active collagen degradation during the development of the zebrafish tail. Developmental Biology, 477, 155–163. 10.1016/j.ydbio.2021.05.016

Yoong, S., O’Connell, B., Soanes, A., Crowhurst, M. O., Lieschke, G. J., & Ward, A. C. (2007). Characterization of the zebrafish matrix metalloproteinase 9 gene and its developmental expression pattern. Gene Expression Patterns, 7(1–2), 39–46. 10.1016/j.modgep.2006.05.005

Zhang, G., Jin, L.-Q., Rodemer, W., Hu, J., Root, Z. D., Medeiros, D. M., & Selzer, M. E. (2022). The Composition and Cellular Sources of CSPGs in the Glial Scar After Spinal Cord Injury in the Lamprey. Frontiers in Molecular Neuroscience, 15, 918871. 10.3389/fnmol.2022.918871

